# Goblet cells mechanically breach the epithelial barrier in gut homeostasis

**DOI:** 10.1101/2025.04.01.646617

**Authors:** Justine Creff, Yuan He, Sandra Bernat-Fabre, Etienne Buscail, Salomé Neuvendel, Vishnu Krishnakumar, Thomas Mangeat, Shi-Lei Xue, Denis Krndija

**Author notes:** These authors contributed equally to this work (Sandra Bernat-Fabre, Etienne Buscail, Salomé Neuvendel).

## Abstract

The intestinal epithelium has to maintain a tight barrier against the harsh luminal environment. Absorptive enterocytes have a polygonal and columnar shape, while mucus-producing goblet cells exhibit a round apical cell shape and a bulky body, raising the question of how epithelial integrity is maintained around these cells. Here, we show that goblet cells induce tight junction fractures between neighboring enterocytes under homeostatic conditions *in vivo*, which are exacerbated during goblet cell hypertrophy, increasing gut permeability. We demonstrate that these fractures arise from a two-component mechanical interaction: goblet cells push and deform adjacent enterocytes, which rupture depending on tissue rheology controlled by myosin II. These findings reveal that the mechanical interplay between goblet cells and neighboring enterocytes is critical for maintaining intestinal epithelial barrier integrity.

## Main Text

Epithelia are specialized tissue barriers, with a critical role in separating internal and external environment and thus maintaining normal tissue function and homeostasis (*1*). The intestinal epithelium functions as a protective barrier throughout adult life, shielding the organism from continuous environmental insults (*2*), including extrinsic mechanical stresses from luminal contents and muscle-driven peristalsis (*3, 4*) as well as intrinsic mechanical stresses due to rapid epithelial turnover (*5*). Adhesive cell junctions – including tight junctions (TJs), adherens junctions (AJs) and desmosomes – enable epithelial barrier, mechanical integrity and collective cell migration (*5*–*7*). TJs are continuous belt-like structures that encircle and seal the paracellular space and regulate the barrier permeability (*8*). AJs, which lie just beneath the TJs, couple the cytoskeletons of neighboring cells by linking transmembrane E-cadherins belts to the cortical actomyosin network, which stabilizes E-cadherin complexes (*9*–*11*). As a consequence, AJs are under tension and transmit subcellular forces exerted by actomyosin networks to the cortex (*12*). Junctional complexes are mechanosensitive and mechanoadaptive, integrating subcellular and tissue-level forces, and modulating cell behavior – e.g., junctional reinforcement upon mechanical load to maintain barrier integrity and tissue cohesion (*10, 11, 13, 14*). Finally, desmosomal plaques are located basolaterally beneath the AJs and are associated to intermediate filaments. They were recently shown to passively respond to extrinsic forces by strain-stiffening, contributing to the mechanical resilience of epithelial tissues (*15, 16*).

The intestinal epithelium consists of at least six different cell types, each with specialized functions and morphologies. Enterocytes, the main cell type, are absorptive cells with a columnar shape and a polygonal apical surface. Mucus-producing goblet cells (GCs), interspersed among enterocytes, are the second most abundant cell type and are characterized by their bulky, round-shaped bodies, due to the presence of large secretory granules (*17*–*19*). While numerous studies have focused on the secretory function of GCs, the impact of their unique geometry and mechanical properties on their integration into the epithelial tissue and the intestinal barrier remains entirely unknown.

Here, we report that under homeostatic conditions *in vivo*, GCs are physically associated with junctional fractures that occur between neighboring enterocytes in the small intestine. Using a combination of 3D tissue imaging, *in vivo* (mouse models), *ex vivo* (explants), and *in vitro* (organoids) force perturbations and theoretical modeling, we found that fracturing arises from the intrinsic mechanical properties and interplay between GCs and adjacent enterocytes. On the one hand, GCs compress and deform adjacent enterocytes, straining their intercellular junctions and promoting fracturing; on the other hand, fracturing depends on global tissue mechanics (rheology), which is controlled by myosin II activity.

### Goblet cells are associated with TJ fractures between adjacent enterocytes under homeostatic conditions

To investigate how GCs are integrated in the small intestinal epithelium, we first assessed TJs by immunofluorescence (ZO-1, occludin) on thick sections of adult murine intestine (jejunum), enabling high-resolution 3D imaging of the villus apical surface *en face* (**Fig. 1A, S1A**). GCs are easily identified due to their round apical shape and absence of an F-actin-rich brush border, in contrast to polygonal enterocytes; their identity was validated by staining for mucus (**Fig. S1B**). We observed that GCs are frequently (>50%) and significantly associated with fractures (**Fig. 1A, S1A**, arrows) in adjacent (1^st^ neighbor) enterocyte-enterocyte (E-E) TJs.

**Fig. 1.**
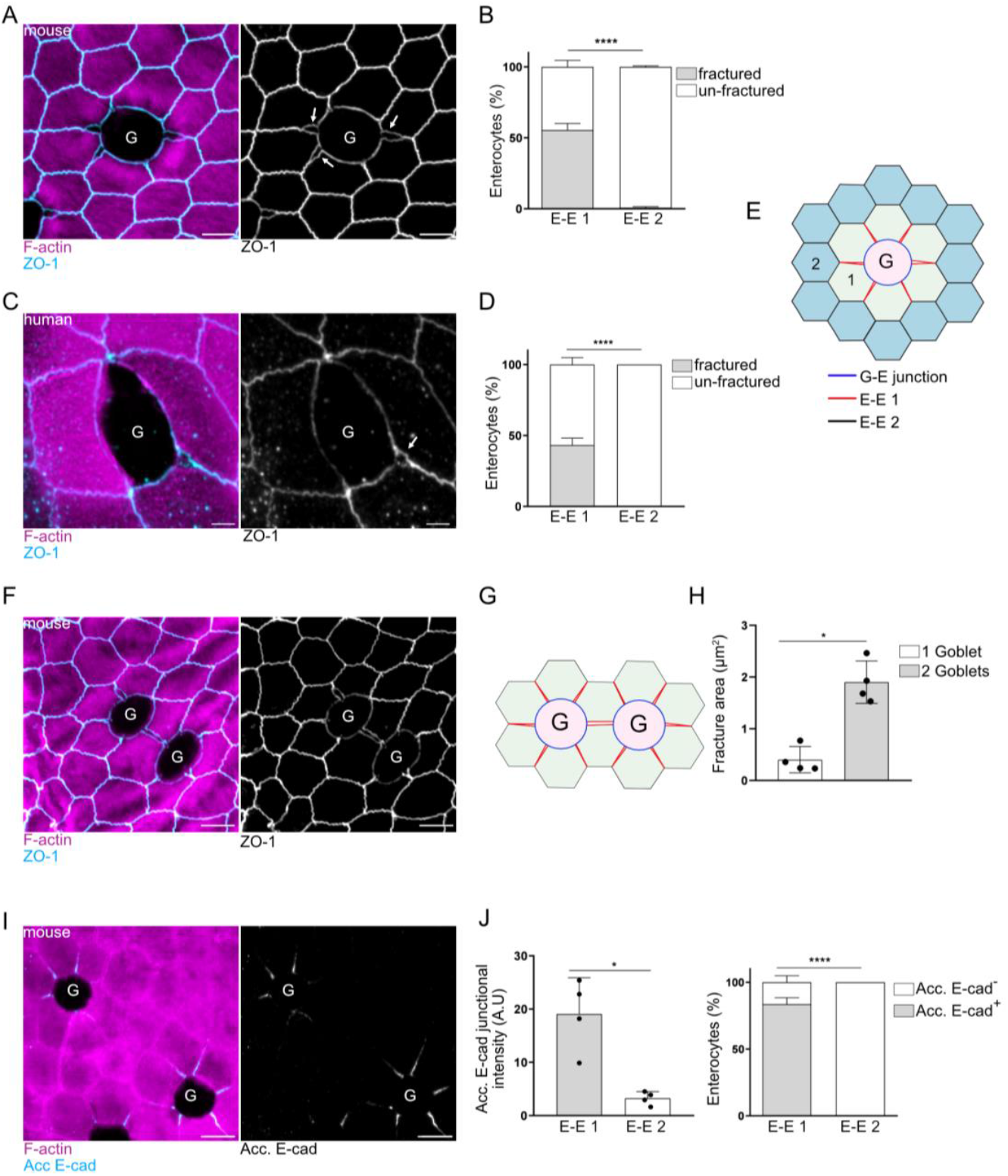
Goblet cells are associated with TJ fractures between adjacent enterocytes under homeostatic conditions. (**A**) Representative *en face* image of murine small intestinal tissue; maximum intensity projection (Z range, 2 µm). White arrows are showing fractures in enterocyte-enterocyte (E-E) 1^st^ neighbor tight junctions (E-E 1). Scale bars, 5 µm. (**B**) Stacked bar graph showing average percentage of enterocytes associated with fractures (GAFs). Multiple t-test, ****: p < 0.0001; n=4 independent experiments. (**C**) Representative *en face* image of thick section of human small intestinal tissue; maximum intensity projection (Z range, 2.2 µm). White arrow is showing fractures in 1^st^ neighbor E-E junctions. Scale bars, 5 µm. (**D**) Stacked bar graph showing average percentage of enterocytes associated with fractures (GAFs) in human tissue. Multiple t-test, ****: p < 0.0001; n=4 independent experiments. (**E**) Schematic representation showing fractures are present in 1^st^ neighbor E-E TJs (E-E 1, red) and absent in G-E (blue) and 2^nd^ neighbor E-E TJs (E-E 2, black). (**F**) Representative image of two GCs in close proximity; maximum intensity projection (Z range, 2.8 µm). ZO-1 (cyan); F-actin (magenta). Scale bars, 5 µm. (**G**) Schematic representation showing the cumulative effect of 2 GCs on fracturing. (**H**) Bar chart displaying average fracture area between 2 GCs. Mann-Whitney test, *: p < 0.05; n=4 experiments. (**I**) Representative image of murine tissue stained for accessible E-cadherin (cyan) and F-actin (magenta); maximum intensity projection (Z range, 3.18 µm). Scale bars, 5 µm. (**J**) Left: Bar chart displaying average accessible E-cadherin junctional intensity; Mann-Whitney test, *: p < 0.05; n=4 experiments. Right: Stacked bar graph showing the average percentage of accessible E-cadherin-positive cells. Multiple t-test, ****: p < 0.0001. (n=3 experiments). Acc. E-cad, accessible E-cadherin.

In contrast, we rarely observed (0.7%) fractures in E-E TJs non-adjacent to GCs (2^nd^ neighbors or farther) (**Fig. 1A, B, E**). The fractures are wider proximal to the GCs, at the goblet-enterocyte-enterocyte (G-E-E) tricellular junctions, and become gradually narrower away from the GC along the E-E bicellular TJs (**Fig. S1C**). In addition, we assessed the presence of fractures at other junctional complexes – at AJ level, fractures were present but ≈3 times less frequent (18%) than at TJs, and no fractures were detected at the level of desmosomes **(Fig. S1D**). Importantly, we also observed fractures in healthy regions of human small intestinal biopsies (ileum) with the same frequency (**Fig. 1C, D**; **Fig. S1E**), suggesting that this phenomenon is conserved between mouse and human.

We asked whether fractures could be the result of extrinsic forces (e.g., luminal shear stress due to digesta flow) or interaction with gut microbiota or immune cells. Remarkably, we found GC-associated fractures in murine embryonic villi (E16.5 embryos, early upon the emergence of villi) (**Fig. S1F**), where digesta, immune system and microbiota are not yet present. To further investigate the influence of extrinsic cues, we turned to murine small intestinal 2D organoids model, which comprises only epithelial cells with functional GCs and mucus, and whose accessible apical surface enables high-resolution imaging of the TJs (**Fig. S2A, B**) (*20*). Similar to what we observed *in vivo*, in the villus-like regions, 50% of goblet cells were associated with TJ fractures (**Fig. S2C-D**). Together, these findings suggest that GC-associated fracturing is epithelium-autonomous, arising from intrinsic properties of the epithelial monolayer. We termed this phenomenon “Goblet cell-Associated Fractures” (GAFs).

Interestingly, we observed an increase in GAF area (five-fold increase) when GCs were in close proximity to each other, i.e., separated by only one row of neighboring cells (**Fig. 1F-H**); this suggests a cumulative effect. To determine whether GAFs represent a breach in the epithelial barrier, we assessed the luminal accessibility of E-cadherin, which is normally inaccessible to antibodies without prior permeabilization. Using a sequential staining approach, tissues were first incubated with an antibody targeting the extracellular domain of E-cadherin before permeabilization, followed by F-actin staining after permeabilization (**Fig. S1G**) (*21*). The “accessible” E-cadherin signal was detected only in GC-adjacent E-E junctions, with over 80% of GCs associated with accessible E-cadherin staining (**Fig. 1I, J**), indicating that TJs around GCs are indeed breached. Taken together, these data suggest that GCs locally disrupt junctional integrity in the gut epithelium under homeostatic conditions.

### Goblet cells mechanically deform the neighboring enterocytes, leading to junctional fractures

Unlike their slender enterocyte neighbors, GCs have a bulky body filled with mucus granules, which deforms the lateral walls of adjacent enterocytes (**Fig. 2A**). Furthermore, super-resolution imaging of desmoplakin, a marker of desmosomal outer dense plaques, revealed a significant and consistent two-fold decrease in intra-desmosomal space at the G-E junctions compared to E-E junctions (**Fig. S3A**), suggesting mechanical compression exerted by goblet cells. Given this, we then explored whether GAF formation is due to the mechanical interactions between GCs and their neighboring enterocytes.

**Fig. 2.**
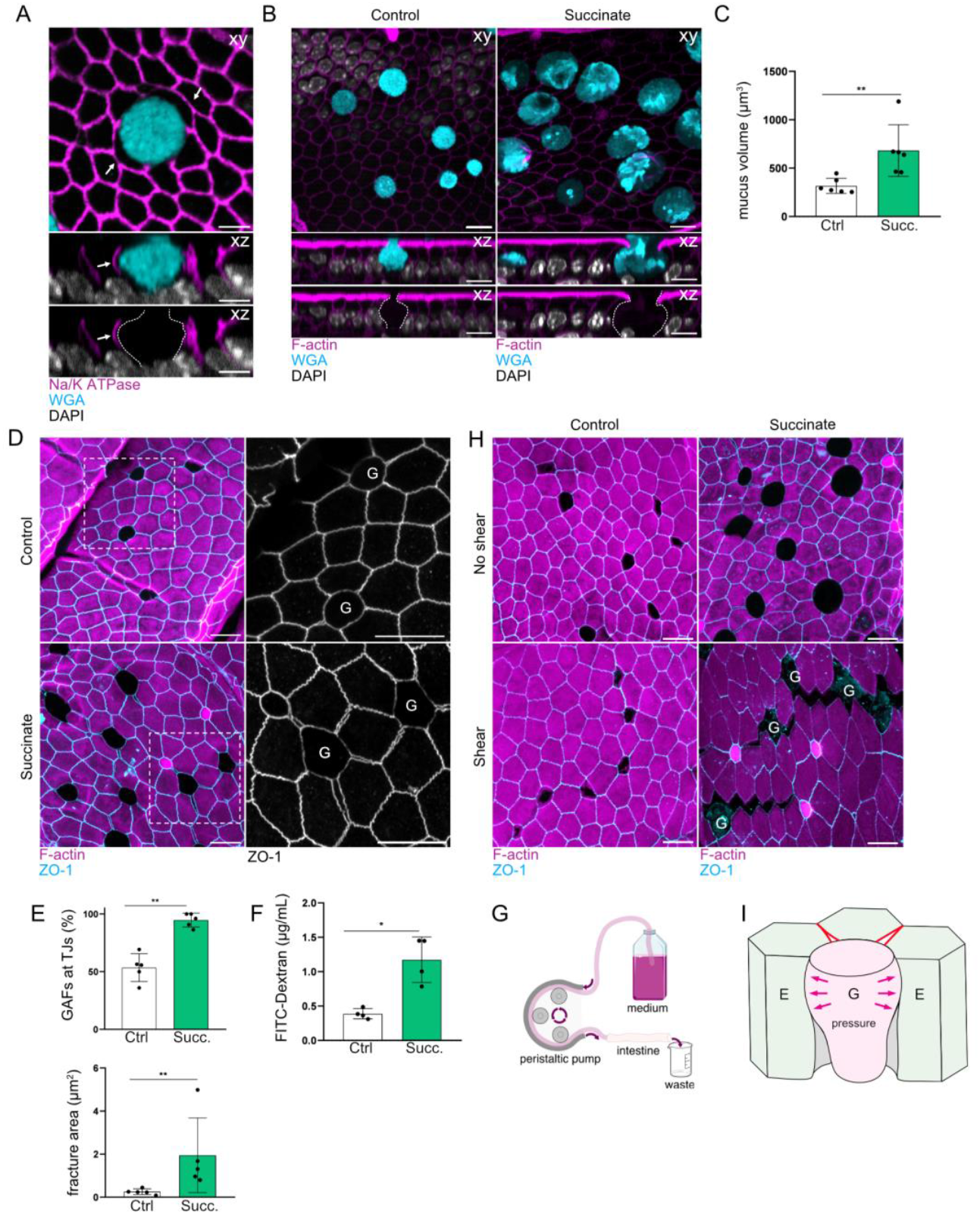
Goblet cells mechanically compress the neighboring enterocytes, leading to their deformation and junctional fractures. (**A**) Representative orthogonal (xy and xz) confocal images from thick intestinal sections. White arrows are showing lateral deformation of GC-adjacent enterocytes. Scale bars, 5 µm. (**B**) Representative xy and xz images of control and succinate tissue showing increased mucus volume, and deformation of goblet-adjacent enterocytes. Scale bars, 10 µm. (**C**) Bar chart displaying average mucus volume. Mann-Whitney test, **: p < 0.01; n=6 experiments. (**D**) Representative images from control and succinate tissues. Boxed regions are shown in higher magnification. Maximum intensity projection (Z range, 4.8 µm, 2.7 µm). Scale bars, 10 µm. (**E**) Top: Bar charts displaying average percentage of GAFs at TJs level. Mann-Whitney test, **: p < 0.01. Bottom: Bar chart showing average fracture area. Mann-Whitney test, **: p < 0.01.; n=5 experiments. (**F**) Bar charts displaying average intestinal permeability. Mann-Whitney test, *: p < 0.05; n=4 experiments. (**G**) Schematic representation of experimental procedure to apply shear stress *ex vivo* on intestinal epithelium. (**H**) Representative images of control and succinate tissues submitted to luminal shear stress; maximum intensity projection (Z range, 3.2 – 4.8µm). Scale bars, 10 µm. (**I**) Schematic representation showing hypothetical pushing forces (pink arrows) emanating from the GC, straining E-E junctions, leading to fracturing. Ctrl, control; Succ., succinate.

From a physical perspective, the stereotypical outward curvature of the G-E junction must arise from GCs having a higher pressure than enterocytes, and such pressure difference should be balanced by an interfacial tensile force at the G-E boundary (see Theory Note for details). The tensile state of the G-E boundary is also evident from its smoothness, which is notably different from the wiggliness displayed by E-E TJs (**Fig. S3B**). Previous studies indicate that the G-E interfacial tension (i.e., the force at the G-E boundary) could act as a “peeling force” on the E-E interface, causing debonding (*22, 23*). Minimal physical modelling of the E-E interface mechanics indeed predicts junctional fracture above a critical G-E interfacial tension (see **Fig. S3C-F** and Theory Note Section 1 for details). Overall, this led us to the hypothesize that GCs exert mechanical forces on their neighbors: the inner pressure within GCs induces a tensile force at the G-E boundary, peeling E-E junctions and causing junctional fracture.

To test this hypothesis directly, we manipulated GC pressure and the corresponding G-E interfacial force by treating mice with succinate, a microbial metabolite which has been proposed to induce GC hypertrophy (an increase in GC mucus volume) and hyperplasia (an increase in the number of GCs) (*24*). We confirmed that the succinate treatment led to a significant increase in GC mucus volume (control: 317.04 µm^3^; succinate: 681.57 µm^3^) (**Fig. 2B, C**), but not GC hyperplasia, as GC numbers remained similar (**Fig. S3G, H**). Hypertrophic GCs were associated with significantly increased fracturing in adjacent E-E tight junctions, showing a three-fold increase in GAF area (control: 0.42 µm^2^; succinate: 1.29 µm^2^) and a nearly two-fold increase in GAF frequency (control: 53%; succinate: 95%) (**Fig. 2D, E**). Despite the increase in fracturing, ZO-1 junctional intensity remained unchanged, suggesting that fractures arise from mechanical perturbations imposed by GC pressure rather than from an underlying structural fragility of the junctions due to decreased ZO-1 (**Fig. S3I**). These results were recapitulated in organoid monolayers, where we induced GC hyperplasia and hypertrophy by inhibiting the Notch and Wnt pathways (*25*) (**Fig. S4**).

Additionally, GC hypertrophy upon succinate treatment resulted in an increase in complete (vertex-to-vertex) GAFs (**Fig. S3J**), consistent with the model’s prediction of complete junctional fracture once the GC pressure threshold is exceeded. Fractures also appeared to extend deeper, as indicated by the increased number of fractures at the level of the AJs and the appearance of fractures at the desmosomes (**Fig. S5**). Importantly, this larger and deeper fracturing was accompanied by a significant increase in intestinal permeability *in vivo* (**Fig. 2F**), suggesting that GC hypertrophy severely weakens epithelial barrier integrity. At the same time, no increase in inflammatory markers or weight loss was observed (**Fig. S6A, B**), suggesting that succinate treatment does not induce immune-mediated tissue damage. The observed increase in fracturing and permeability is therefore likely to be driven by epithelial mechanics rather than disease. To test whether increased fracturing observed with GC hypertrophy further compromises tissue integrity under extrinsic mechanical stress, we applied a defined flow through the intestinal lumen *ex vivo* using a peristaltic pump system (shear stress estimated at 1Pa) (**Fig. 2G**). This shear stress challenge resulted in a striking increase in fractures emanating from GCs in tissues from succinate-treated mice, with fractures extending further (≥3 neighboring cells) compared to control tissue, which remained unaffected (**Fig. 2H, S6C**). Taken together, these data suggest that GCs exert pressure on neighboring enterocytes, leading to junctional ruptures (**Fig. 2I**), which can be further exacerbated by increased GC volume (goblet hypertrophy). Importantly, we find that this leads to functional consequences, notably an increase in intestinal permeability and heightened susceptibility to mechanical stress from the environment.

### Active response of the tissue to goblet cell pressure

Cellular junctions are mechanoresponsive. This response involves different junctional components at different levels (*6, 26*). E-cadherin, a key marker of AJs, is known to respond to mechanical stress by clustering (*11*). Interestingly, we found a striking increase in E-cadherin junctional levels in GC-adjacent E-E AJs compared to non-adjacent AJs (**Fig. 3A, B**). Moreover, E-cadherin signal levels were heterogeneous along the junctions, progressively increasing towards the goblet cell (**Fig. 3C**), a pattern also observed in 2D organoids (**Fig. S7**). This accumulation prompted us to investigate whether it indicated higher strain near GCs. To test this, we analyzed the distribution of tricellulin, a marker of tricellular junctions (TCJs), which are associated with tension hotspots and mechanical failure points (*27, 28*). Notably, tricellulin levels were elevated at G-E-E TCJs compared to E-E-E TCJs (**Fig. 3D, E**). Moreover, the tricellulin signal at the GAFs was disorganized, exhibiting multiple puncta, in contrast to the single puncta observed at unfractured TCJs (**Fig. 3D, F**), supporting the hypothesis that fractures initiate at the G-E-E TCJ and propagate towards the E-E bicellular junction (**Fig. S1C**). At the level of desmosomes, we found a 2.8-fold increase in desmocollin – a desmosomal cadherin (*29*) – in GC-adjacent E-E desmosomes (**Fig. 3G, H**), suggesting local mechanoadaptation of desmosomes. Altogether, our results show that junctional protein accumulation at TCJs, AJs and desmosomes in enterocytes correlates with their proximity to GCs. We propose that the tissue actively responds by mechanoadaptation to mitigate the stresses arising from goblet cell compression and fracturing.

**Fig. 3.**
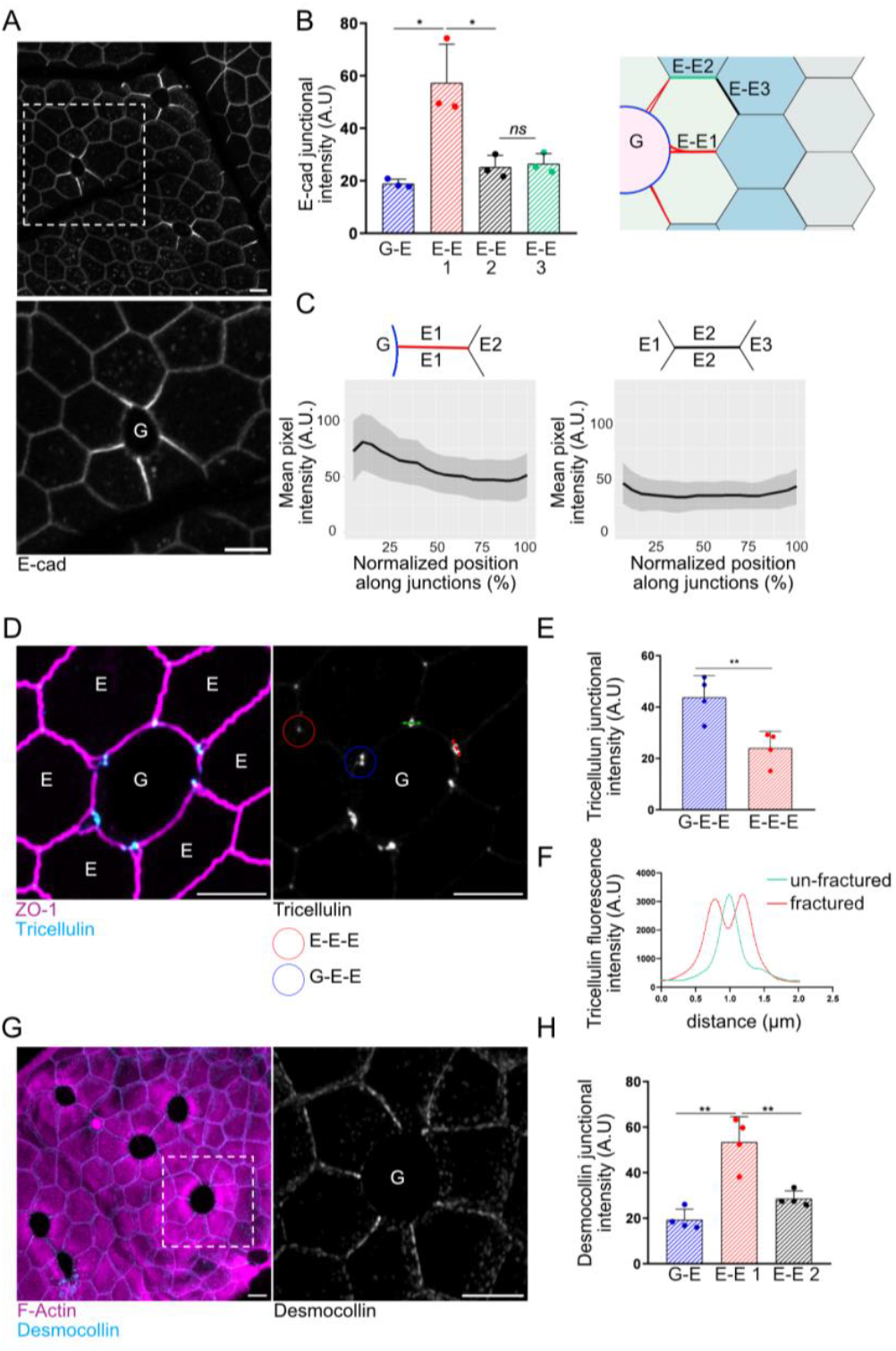
Active mechanoadaptation of the tissue to goblet cell pressure. (**A**) Representative *en face* image; boxed region is shown in higher magnification; maximum intensity projection (Z range, 1.8 µm). Scale bars, 5 µm. (**B**) Bar chart showing average junctional E-cadherin intensity. Multiple t-test, ns: non-significant; *: p < 0.05; n=3 experiments. (**C**) Average line intensity profiles for junctional E-cadherin E-E 1^st^ neighbor (left) and E-E 2^nd^ neighbor (right) along rescaled E-E junctions, as indicated. (**D**) Representative *en face* image; G-E-E are encircled in blue and E-E-E in red; maximum intensity projection (Z range, 1.4 µm). Scale bars, 5 µm. (**E**) Bar charts showing average tricellulin intensity in G-E-E or E-E-E TCJs. Unpaired t-test. *: p < 0.05; n=4 experiments. (**F**) Line intensity profiles for tricellulin at fractured (red) and un-fractured (green) tri-cellular junctions; these profiles correspond to lines shown in E. (**G**) Representative *en face* image; maximum intensity projection (Z range, 1.5 µm). Boxed region is shown in higher magnification. Scale bars, 5 µm. (**H**) Bar chart displaying average desmocollin junctional intensity. Multiple t-test, **: p < 0.01; n=4 experiments.

### Goblet cell-associated fracturing depends on mechanical properties of neighboring cells

Because junctional fracturing often correlates with locally increased active stresses generated by actomyosin contractility (*30*), and given the increased E-cadherin signal we observed (**Fig. 3A**), indicative of local mechanical strain, we next investigated whether and how tissue mechanics contribute to GAFs.

To determine whether there are differences in contractility between GCs and their neighboring enterocytes, we first assessed the levels of non-muscle myosin IIA (NMIIA) in the apical actomyosin belts using live imaging of gut explants derived from mice expressing GFP-tagged endogenous NMIIA (*31*). We found that GCs exhibited higher NMIIA levels in their apical belts compared to the myosin belts of enterocytes (**Fig. 4A, B**). However, phospho-myosin light chain (T18/S19) (P-MLC) staining, a marker of active myosin, was not detectable in the apical belts of GCs compared to the junctions of neighboring enterocytes (**Fig. 4C, D**), suggesting lower contractility. To confirm these results and to assess the tensile forces, we performed laser cuts of the individual G-E or surrounding E-E AJ junctions *ex vivo*, in gut explants (*5*) (**Fig. S8**). The recoil velocity was 4-fold higher at the E-E junctions than at the G-E junctions, indicating that the GC myosin ring does not confer increased contractility. Beyond its role in contractility, myosin II crosslinking of cortical actin filaments is critical for the mechanical properties of cells and junctions (e.g., rigidity), which influence cell shape and junctional strength (*32*–*34*). We hypothesized that epithelial myosin II could therefore significantly influence the formation of GAFs. We tested this using an inducible, intestinal epithelium-specific knockout mouse model of NMIIA (NMIIA^KO^; **Fig. 4E, S9A**) (*35*). We found that NMIIA depletion resulted in a significant reduction in GAF area (control: 0.64 µm^2^; NMIIA^KO^: 0.33 µm^2^) (**Fig. 4E, F**), and a decrease in E-cadherin enrichment (**Fig. S9B, C**), while no change in GC mucus volume was detected (**Fig. S9D**).

**Fig. 4.**
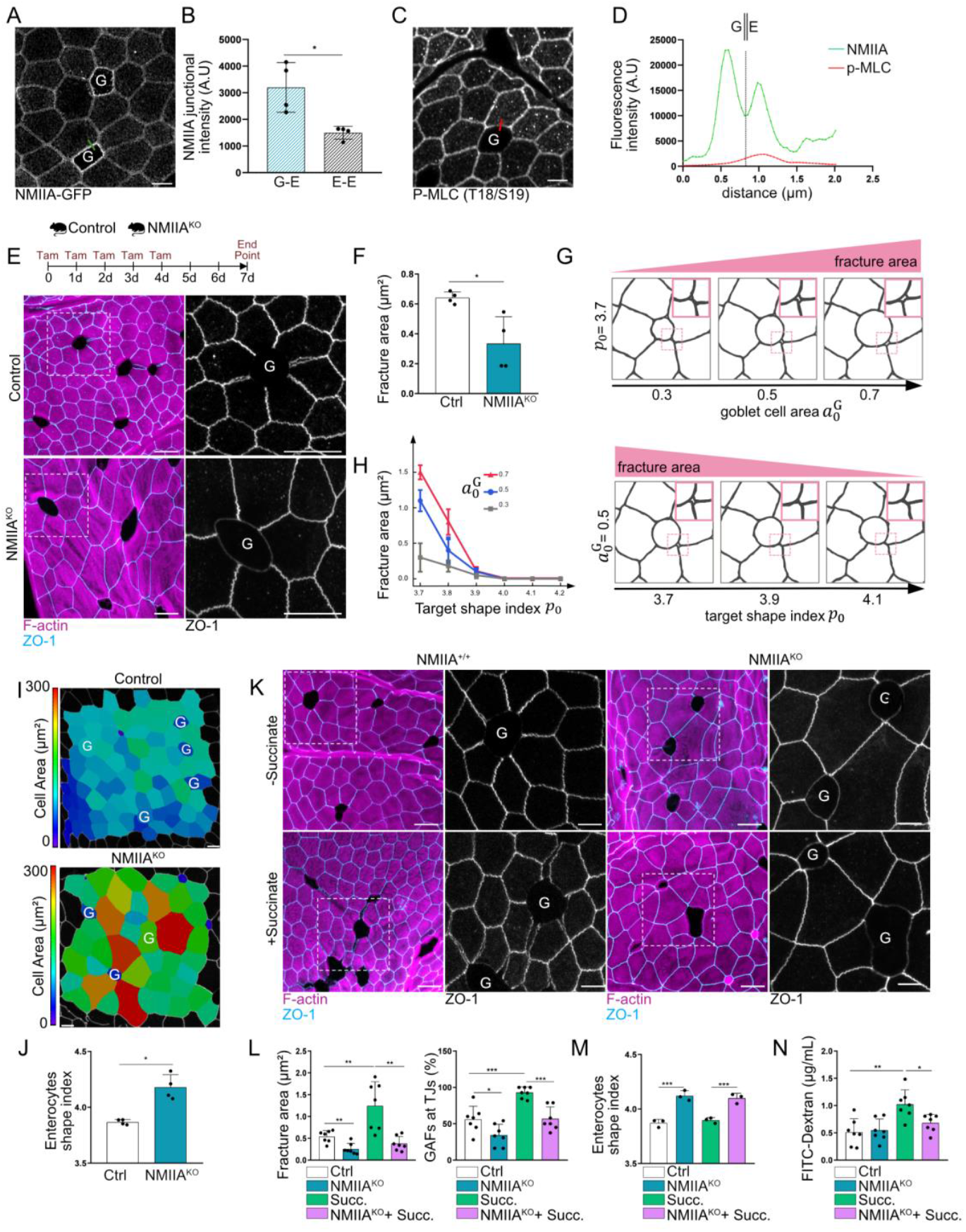
Goblet cell-associated fracturing depends on global tissue mechanical properties. (**A**) Representative confocal image of live intestinal tissue showing apical NMIIA-GFP signal. Maximum intensity projection (Z range, 1.2 µm). Scale bar, 5 µm. (**B**) Bar chart displaying average NMIIA junctional intensity; maximum intensity projection (Z range, 2 µm). Mann-Whitney test, *: p < 0.05; n=4 experiments. (**C**) Representative image showing apical pMLC signal. Scale bar, 5 µm. (**D**) Line intensity profiles of NMIIA-GFP and phospho-MLC corresponding to lines shown in images A and C. (**E**) Top: Schematic representation of experimental procedure for NMIIA^KO^ mice; Tam, tamoxifen. Bottom: Representative images from control and NMIIA^KO^ mice; maximum intensity projection (Z range, 4.5 µm and 3.8 µm). Boxed regions are shown in higher magnification. Scale bars, 10 µm. (**F**) Bar chart showing average fracture area. Mann-Whitney test. *: p < 0.05; n=4 experiments. (**G**) Simulations of the tissue mechanics model with two main parameters: the goblet cell area 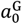 and the shape index of enterocytes *p*_0_ (relevant to tissue rheology). The model predicts that GAFs increase with the goblet cell area 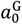 (cell shape index p0 =3.7, top panel), but decrease with the shape index *p*_0_ (goblet cell area 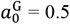, bottom panel). (**H**) Predicted fracture area as a function of goblet cell area 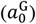 and shape index of enterocytes (*p*_0_). (**I**) Representative heatmap images displaying apical cell area in control and NMIIA^KO^ mice. Scale bars, 5 µm. (**J**) Bar chart displaying average enterocyte shape index. Mann-Whitney test. *: p < 0.05; n=4 experiments. (**K**) Representative *en face* images from control (NMIIA^+/+^) and NMIIA^KO^ mice, with and without succinate treatment. Scale bars, 10 µm. Boxed regions are shown in higher magnification. Maximum intensity projection (Z range, 3.0 – 4.5 µm). Scale bars, 5 µm. (**L**) Bar charts displaying average fracture area (left) and average percentage of GAFs (right), according to the conditions. Multiple t-test, *ns*: non-significant, *: p<0.05, **: p < 0.01, ***: p<0.001, ****<0.0001; n=7 experiments. (**M**) Bar chart displaying average enterocyte shape index. Multiple t-test. ***: p < 0.005; n=3 experiments. (**N**) Bar chart showing average intestinal permeability. Multiple t-test; *: p<0.05, **: p < 0.01; n= 7 experiments. Succ., succinate.

This result, however, contradicted the peeling model for E-E interfaces (**Fig. S3C**), which predicted that a decrease in junction rigidity would facilitate fracture as junctions become more susceptible to GC-induced deformation (see **Fig. S3E, F** and Theory Note Section 1 for discussion). A possible explanation for this discrepancy is that the peeling model considers only a single E–E junction, whereas tissue-level models emphasize how global cell shape and packing influence epithelial mechanics. After NMIIA depletion, enterocytes no longer formed regular pentagonal or hexagonal arrays but became floppy and disorganized (**Fig. 4E, I**). Given previous findings that cell shape tightly correlates with tissue rheology (*36*–*38*), we reasoned that junctional fracture may not be solely a local mechanical event, but rather depend on mechanical properties at the tissue level. Recent theoretical work incorporating fracture into vertex models further supports the idea that changes in tissue rigidity modulate susceptibility to local fractures (*39, 40*). Thus, if GC-adjacent enterocytes rearrange to make the tissue more fluid-like, they may dissipate GC-induced forces and thereby alleviate GAFs.

To explore this hypothesis, we developed an extended vertex model that retained the key physical components of the classical vertex model (*37*–*42*) – namely, the existence of a preferred cell shape and area – while allowing for complex junctional deformations and ruptures, which were crucial for modeling GC and GAF shapes (for more details, see Theory Note Section 2 and **Fig. S10A**). We found that the presence of GAFs was governed by two key parameters (**Fig. 4G**): GC size/pressure 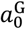, as in the previous model, and tissue rheology, which can be inferred from the “cell shape index” *p*_0_ (i.e., the cellular perimeter-to-area ratio) (*36*–*38*). As the shape index increases, the tissue becomes more fluid-like (*41*), which prevents the formation of GAFs (**Fig. 4G, H**). This leads to the prediction that NMIIA^KO^ enterocytes should be more fluid, i.e., have a higher shape index. We tested this by segmenting control and NMIIA^KO^ tissue, finding increased apical cell area and an increased cell shape index for the KO (control: 3.86; NMIIA^KO^: 4.18) (**Fig. 4I, J; Fig. S9E**), thus confirming our prediction. Together, these results support our hypothesis that fracturing depends not only on goblet cell pressure, but also on overall cell shape and mechanical properties regulated by myosin II activity.

To further test model predictions and determine whether myosin II depletion could prevent the increased fracturing and barrier breaching observed with GC volume increase (**Fig. 2**), we induced GC hypertrophy *in vivo* in NMIIA^KO^ mice and in blebbistatin-treated 2D organoids (**Fig. S11A, S12A**). Hypertrophy-induced GAFs were significantly reduced both *in vivo* in NMIIA^KO^ and *in vitro* (**Fig. 4K, L, S12B-D**), including deeper fractures at AJs and desmosomes (**Fig. S11B-E**). Consistent with our previous findings, NMIIA^KO^ enterocytes, treated or not with succinate, displayed an increased cell shape index compared to control or succinate-treated epithelium (**Fig. 4M**), further emphasizing the importance of enterocyte rheology in the fracturing process. Importantly, the GC hypertrophy-associated increase in intestinal permeability observed *in vivo* was significantly reduced in NMIIA^KO^ mice (**Fig. 4N**). Overall, our results indicate that GAF formation depends on both goblet cells and their immediate neighbors – a two-component system in which the goblet cell body pushes and deforms neighboring enterocytes, straining and fracturing their junctions, which depends on tissue rheology controlled by myosin II.

## Discussion

Here we show that under homeostatic conditions *in vivo*, intestinal goblet cells (GCs) exert local mechanical compression on adjacent enterocytes due to their bulky shape and volumetric expansion, resulting in fracture of junctions between enterocytes, as a function of enterocyte mechanics controlled by myosin II.

The integrity of the intestinal epithelial barrier is critical for maintaining homeostasis, and its disruption can trigger pathologies such as inflammatory diseases (IBD) and tumorigenesis (*43*). The barrier function of the intestinal epithelium is maintained by adhesive junctional complexes, particularly TJs, which are the primary physical determinant of the epithelial barrier (*8*), assumed to form a continuous, uninterrupted seal between adjacent cells, blocking the passage of bacterial toxins and pathogens into the underlying tissues (*43*). However, most studies of epithelial cell junctions have focused predominantly on cell lines or single “main” cell types, such as enterocytes, using thin histology sections. In contrast, employing thick tissue sections and high-resolution 3D confocal microscopy, we found that approximately 50% of GCs are associated with fractures at the TJ level. This GC-associated junctional breaching is even more prevalent than what we can detect by imaging TJ markers, as revealed by the E-cadherin accessibility assay, which showed 80% of GC-adjacent enterocytes with breached TJs. Junctional fracturing in epithelial tissues has previously been observed in cultured monolayers subjected to extrinsic stretch (*15*), in placozoans under high motile stress (*40*), and during hydraulic fracturing of blastocysts (*44*). However, goblet cell-associated fractures represent the first example of an adult epithelium undergoing self-induced rupture under homeostatic conditions.

While the epithelial integrity and mechano-responsiveness of individual junction types have been relatively well studied, their comparative adhesive strength and resistance to mechanical stress *in vivo* remain unclear. Interestingly, we found that TJs fracture more frequently (50%) than AJs (18%), while no fractures were observed at the desmosome level under homeostatic conditions. This suggests that TJs are more fragile and less resistant to mechanical stress than the underlying junctions. Notably, fractures predominantly originate at tricellular junctions (TCJs) at the G-E-E interface, before propagating along the E-E bicellular junction. These results are consistent with observations that TCJs are points of high tension in the tissue and may represent “weak spots” in the epithelium that are more prone to fracturing (*27*). Remarkably, we found that enterocytes accumulate transmembrane proteins E-cadherin and desmocollin in response to the presence of GCs, suggesting junctional reinforcement, as previously shown *in vitro* – reviewed in (*11*). This accumulation of junctional proteins may serve to increase junctional strength and prevent further rupture.

Junctional reinforcement has previously been described as a response to increased tension generated by intrinsic cellular contractility in developing embryos and cultured epithelial monolayers – reviewed in (*14*), suggesting that GC contractility may play a role. However, the prominent apical myosin ring in GCs does not appear to be contractile, leading us to speculate that it has a more passive, scaffolding role in containing mucus granules within the cell. In addition, we found that GAFs depend on global tissue mechanical properties (i.e., solidity/fluidity), which we manipulated by perturbing myosin II. Reduced myosin expression or activity resulted in a more fluid tissue that was less prone to rupture. Such mechanical fluidization could act by preventing stress accumulation in the tissue by allowing the cells to adjust their shape/size and position and thus adapt to abnormal mechanical forces, protecting the tissue from physical damage. These findings are in line with recent *in vitro* experimental work (*15*) and theoretical studies of rupture in epithelia (*39*), which show that myosin and the associated solid- or fluid-like state of the tissue play a critical role in fracturing processes.

The presence of GCs is necessary for fractures to occur, as no fractures were detected in regions within the epithelium without GCs. Our results indicate that fractures are promoted by the volume of GCs and the pressure they exert on neighboring enterocytes. In our study, succinate treatment was used to mimic a physiological response to parasitic infection, a condition previously described as inducing GC hyperplasia and hypertrophy, leading to an increase in GC numbers and mucus hypersecretion. GC hyperplasia and hypertrophy was shown to promote parasite clearance through the “weep and sweep” response and thus “protect” the tissue (*24, 45*). While we did not observe GC hyperplasia in our experiments, possibly due to technical reasons, we observed prominent GC hypertrophy, characterized by a marked increase in intracellular mucus volume. Interestingly, GC hypertrophy was closely associated with a significant increase in fracture size (both area and depth) and number, ultimately leading to increased intestinal permeability, challenging the widely held view that GC hypertrophy is solely protective.

Although increased permeability was observed following succinate treatment, no evidence of inflammation was detected. This is somewhat unexpected, as elevated paracellular permeability is generally associated with enhanced bacterial translocation and inflammatory responses. We hypothesize that in pathological contexts of active infection, pathogen-induced GC hypertrophy may facilitate the translocation of microbes and luminal toxins, ultimately triggering inflammation. Conversely, under homeostatic conditions, compensatory mechanisms such as mucus hypersecretion may counterbalance GC-induced junctional disruption by enhancing microbial entrapment and thereby limiting exposure of the underlying epithelium. However, when luminal shear stress was applied *ex vivo*, GC hypertrophy severely compromised tissue integrity, creating large intercellular gaps that spanned multiple rows of adjacent cells. Shear stress is physiologically relevant, as it can increase in conditions such as partial intestinal obstruction or luminal narrowing caused by inflammation, fibrosis, strictures, or tumors, where fluid velocity increases. These findings suggest that an increase in GC volume (hypertrophy) is necessary and sufficient to increase fracturing and weaken the intestinal epithelium, making it more vulnerable to mechanical insults, which could further exacerbate damage and trigger inflammation.

GCs have been identified as entry points for *Listeria monocytogenes* invasion and infection (*21*), a process that requires luminally accessible E-cadherin. This is consistent with our findings of GAFs and the associated barrier breach, which exposes E-cadherin and could thus facilitate bacterial adhesion and tissue infection. GCs have also been identified as mediators of interactions between the immune system and luminal content through transcytosis-based pathways – the goblet cell-associated antigen passages (GAPs) – that enable the sampling of luminal antigens and their delivery to underlying immune cells (*46*). The junctional fractures (GAFs) described here may have a similar function, physically facilitating antigen sampling by immune cells. However, we have also observed GAFs in the embryonic mouse intestine, where the lumen is empty, and neither the immune system nor the microbiota are present. Remarkably, GAFs were observed in both mouse and human intestinal tissue, suggesting it may be conserved across species.

What is the possible role of GAFs? Fracturing near the goblet–enterocyte interface may serve as a local mechanical dissipation mechanism – a controlled failure to mitigate the risk of larger-scale epithelial disruption. This concept parallels developmental processes, where local deformation and cellular rearrangement dissipate mechanical forces during morphogenesis (*7, 9, 38, 47, 48*). Consistently, we observed an active tissue response to GC compression, characterized by mechano-sensitive reinforcement of junctional adhesion in GC neighbors. Analogous stress-relief phenomena have been observed in a few other biological systems, such as the migratory fracturing in Trichoplax adhaerens (*40*), where epithelial rupture is thought to prevent tissue failure under motile stress. However, this compensatory response appears insufficient under conditions of GC hypertrophy, where elevated compressive forces exacerbate fracturing and compromise barrier integrity – highlighting the limits of epithelial plasticity under sustained mechanical stress.

Our unexpected results demonstrate that goblet cells challenge the intestinal epithelial barrier by creating structural vulnerabilities that may contribute to intestinal pathologies, including infection, inflammation and disease progression. In conclusion, we show that the mechanical interplay between goblet cells and neighboring enterocytes promotes junctional fractures, compromising the epithelial barrier. We propose that the mechanical properties of individual cells – both goblet cells and enterocytes – are key factors in maintaining junctional integrity and preserving the intestinal epithelial barrier.

## Acknowledgments

We thank E. Hannezo, M. Suzanne, B. Bénazéraf, E. Theveneau, A. Davy, R. Galupa, and all members of the Krndija lab for their critical feedback and valuable suggestions. We acknowledge the help and contribution of the ANEXPLO mouse facility and Light Imaging Toulouse CBI (LITC) facility at the CBI, and TRI-IPBS imaging facility (supported by the French National Research Agency – ANR-24-INBS-00005 FBI Biogen), members of the national infrastructure France-BioImaging. Special thanks to Laurent Malaquin (LAAS) for luminal shear stress simulations, Alexandre Lucas (We-Met platform, I2MC, Toulouse) for conducting the ELISA tests, and Silvia Kočanová for assistance in acquiring human samples.

## Author contributions

Conceptualization: D.K.; Methodology: J.C., Y.H., S.-L.X., D.K.; Investigation: J.C., S.N., S.B.F., V.K., T.M., S.-L.X., E.B.; Project administration: J.C., D.K.; Supervision: S.-L.X., D.K.; Funding acquisition: D.K.; Writing – original draft: J.C., D.K.; Writing – review & editing: J.C., D.K.

## Data and materials availability

All data are available in the main text or the supplementary materials.

