## Supplementary materials for "Goblet cells mechanically breach the epithelial barrier in gut homeostasis"


**The PDF file includes:**

Materials and Methods
Theory Note
Figs. S1 to S12
Supplementary references

### Materials and Methods

Mice

All mouse husbandry and handling procedures were approved and conducted in accordance with European and national regulation (Protocol Authorization APAFIS #38994-2023062718345638). All mice were kept in the Specific and Opportunistic Pathogen-Free (SOPF) animal facility at the CBI. Comparisons were made between age-matched littermates (8–12 weeks old). As no phenotypic differences were observed between males and females, mice of both sexes were used in all experiments. Villin:Cre$^{ERT2}$ mice (*49*) were crossed with Myh9$^{flox}$ (*35*), obtained from the Mutant Mouse Resource and Research Center (MMRRC; stock 032096-UNC), to generate NMIIA$^{KO}$ mice. Villin:Cre$^{ERT2}$ and NMIIA$^{GFP}$ (*31*) mice were provided by Dr. D. Matić Vignjević (Institut Curie).

*Tamoxifen and drug treatments*

To induce intestinal epithelium-specific Myh9 (NMIIA) knockout, Villin:Cre$^{ERT2}$/Myh9$^{fl/fl}$ and Myh9$^{fl/fl}$ (control) mice were intraperitoneally injected with 100 µL of tamoxifen (50 mg/kg; Euromedex SE-S1238) once a day for 5 consecutive days and culled 3 days after the last injection, as indicated. For succinate treatment, mice received 100 mM succinate (Sigma, S9637) in drinking water for 7 days (*24*), as indicated. For DSS treatment, mice received 3% DSS (Sigma, 42867) in drinking water for 7 days, after which they were culled.

Antibodies

For immunofluorescence, we used the following antibodies: rat anti-ZO-1 (Millipore, MABT11) at 1/200, rat anti-E-cadherin (Invitrogen, 131900) at 1/200, rabbit anti-occludin (Proteintech, 27260-1-AP) at 1/50, sheep anti-desmocollin-2 (Biotechne, AF7490) at 1/100, Rabbit anti-tricellulin (Invitrogen, 48-8400) at 1/50, mouse anti-desmoplakin (Progen, 651109) at 1/50, rabbit anti-phospho-Myosin Light Chain (P-MLC) (T18/S19) (Cell Signaling, 3674S) at 1/50, rabbit anti-Na/K ATPase (abcam, ab-76020) at 1/400. For human tissue staining, mouse anti-ZO-1 (Invitrogen, 1A12) at 1/100 dilution was used. All secondary antibodies were purchased from Molecular probes and used at a dilution of 1/200. F-actin was stained using rhodamine- or Alexa-647-conjugated phalloidin (Invitrogen R415, A30107) at 1/200 dilution. Intracellular mucus was stained using WGA-rhodamine (VectorLabs, RL-1022) at 1/100 dilution. DNA was visualized using DAPI (Sigma, D9542) at 5µg/mL. For western blotting, we used: rabbit anti-NMIIA (Biolegend, 909801) at 1/5000 and rabbit anti-GAPDH (Sigma, G9545) at 1/10000, followed by horseradish peroxidase-conjugated anti-rabbit (Invitrogen, G21234) as a secondary antibody, at 1/10000 dilution.

3D immunofluorescence staining

Mice were culled as indicated, and the small intestine (jejunum) was isolated, flushed with PBS, cut into 3-5 cm fragments, and fixed in 4% paraformaldehyde (Electron Microscopy Sciences) in PBS for at least 1 h at 37°C, followed by three washes in PBS. Fixed tissue was hand cut with a scalpel and permeabilized with 1% Triton X-100 (Sigma) in PBS for 1h at RT. Slices were then incubated with primary antibody in PBS overnight at RT under agitation. After three 1-hour washes in PBS, secondary antibody was added along with DAPI, and, optionally WGA and phalloidin, and incubated overnight at RT. Samples were washed three times for 1h each and mounted on slides using mounting medium (Aqua-Poly/Mount, Polysciences), according to the manufacturer's instructions.

<u>Human tissue staining</u>

Human intestinal biopsies were obtained from the CHU Rangueil in Toulouse with informed consent from each patient (MTA 24 134C). Patients were diagnosed with inflammatory pathologies, and biopsies of ileum were taken from non-inflamed regions. Tissues were fixed in 4% paraformaldehyde O/N at RT, and washed three time with PBS. Sections were hand-cut with a scalpel and slices were then permeabilized with 1% Triton X-100 in PBS for 2 h at RT and stained as described above.

<u>Luminal shear stress</u>

*Ex vivo perfusion*

Mice were sacrificed and the jejunum portion of the small intestine was collected and cut into ≈5 cm segments. Each intestinal segment was mounted onto a peristaltic pump system via a gavage needle, allowing controlled luminal perfusion. Segments were perfused with Leibovitz's L-15 medium (1145056, Gibco) at a constant flow rate of 23 mL/min for 5 min. Tissues were then fixed and stained as described above.

*Simulation of luminal shear stress*

Numerical simulations of luminal shear stress were performed using COMSOL Multiphysics and finite element analysis (FEA). The Laminar Flow physics module was used to solve the incompressible Navier-Stokes equations under steady-state conditions. A 2D/3D geometry representative of a murine intestinal segment was constructed to model the internal topography of the jejunum. The model consisted of a 10 mm-long intestinal tube containing 260 villi (500 µm in height), homogeneously distributed along the luminal surface. The working fluid was defined with a density of 1 g/mL and a dynamic viscosity of 10 mPa·s, approximating the properties of the perfusion medium. A volumetric flow rate of 23 mL/min was imposed at the inlet, assuming a uniform velocity profile. Boundary conditions included no-slip at all solid surfaces and zero pressure at the outlet. Shear stress values were extracted from the computed stationary solution.

<u>Laser ablations of cell junctions</u>

Laser cuts of cell-cell junctions were performed as previously described (*5*). Briefly, NMIIA$^{GFP}$ mice (*31*), where non-muscle myosin heavy chain II-A is endogenously tagged with GFP, allowing visualization of adherens junction-associated actomyosin, were used to conduct junctional laser cuts. Living intestinal tissue slices were hand-cut using a scalpel and placed into a 35-mm glass-bottom culture dish (Fluorodish) containing 200 µL of Leibovitz medium. A slice anchor (SHD-26GH/10; Harvard Apparatus) was placed on top of the tissue to minimize sample drift. To ensure consistency and maintain tissue viability and tension, imaging and laser ablations were conducted within 45 minutes following the dissection. The intestinal tissue was imaged using a two-photon laser-scanning microscope (LSM710, Carl Zeiss) in single-photon mode (488 line), at a resolution of 512 × 512 pixels (pixel size = 0.09 µm) with a bidirectional scan lasting 1 s. Cell junctions were ablated using a Chameleon Vision II laser set at 800 nm (three iterations), with laser power of 1-2 mW. The iteration number was optimized to efficiently ablate cell junctions without causing cavitations or additional cell damage. As tissue recoil typically plateaued around 50 s, images were acquired at 1 s intervals (δt =1 s) for a total of 60 s. Analysis were performed using a custom ImageJ macro as previously described (*50*). Briefly, images were denoised using a filtering process to improve the signal-to-noise ratio. Recoil speed was determined by measuring the vertex-vertex distance $d_{vv}$ at each time frame, with each value subtracted from $d_{vv}(t_0)$ to obtain the amount of strain or recoil (ε(t)) at each time point. As junctional strain exhibits a single exponential growth with a defined plateau after ablation, it can be modeled as a Kelvin-Voigt fiber, by fitting to the following equation $\varepsilon(t) = \frac{F0}{E} \mathrm{x}(1 - e^{-[(\frac{E}{\mu})\mathrm{x}t]})$, where F0 is the tensile force at the junction before ablation, E is the elasticity of the junction, and µ is the viscosity coefficient. The initial recoil parameter $\frac{F0}{\mu}$ can be extracted after fitting. The fitting and plotting of ε(t) and initial recoil values was done in GraphPad Prism.

Intestinal permeability assay

Mice were orally administered 4 kDa FITC-Dextran (Sigma, 46944) at 8 mg per mouse in a randomized order. Two hours later, mice were anesthetized using isoflurane, and blood was collected by cardiac puncture and allowed to coagulate for 30 min at RT. Blood samples were centrifuged (2200 x g, 15 min), and serum was collected. Serum from non-gavaged mice was used as a blank. FITC-Dextran concentration in serum was measured in duplicates using a Fluoroskan FL with SkanIt software plate reader.

Cytokine quantification by ELISA

Cytokine levels (TNF-α, IL-6, and IFN-γ) were measured using the ELLA automated ELISA system (Biotechne). Control, succinate or DSS treated mice (used as a positive control) were anesthetized using isoflurane and blood was collected by cardiac puncture. Blood samples were centrifuged (1000 x g, 20 min) and serum was collected and diluted according to the manufacturer's protocol.

Organoids

*Intestinal crypt isolation and 3D organoid culture*

Wild type mice (2 months old) were sacrificed and the jejunum was isolated, cut longitudinally, washed in PBS and minced with scalpel. Tissue was dissociated in PBS-EDTA 2mM solution for 30min at 4°C. After shaking, the fraction containing crypts was filtered through a 70 µm strainer (Corning). The solution was centrifuged at 1200 rpm at 4°C for 4 min to separate crypts from single cells. The supernatant was removed and the pellet was resuspended in 10 mL of resuspension solution (DMEM/F12 + 2% AA) and filtered through a 70 µm strainer to remove the

villi. The solution was then centrifuged at 1200 rpm at 4°C for 6 min. The pellet was resuspended in 1/1 Matrigel (Corning, 356231)/ENR medium and plated as 40µL drops in 24-wells plates. After incubation for 15 min at 37°C and 5% $CO_2$ in a humidified atmosphere, the wells were filled with 350 µL of ENR medium. Organoids were passaged every 3-4 days by mechanical dissociation.
ENR medium: Advanced DMEM F12 (Gibco, 12634010), supplemented with 2% AA (Gibco, 11570486), 2.5% Glutamax (Gibco, 13462629), 100 ng/mL Noggin (StemCell Technologies), 20 ng/mL EGF (Peprotech, 315-09), 500 ng/mL R-Spondin 1 (Peprotech, 315-32), 10 ng/mL mbFGF (Peprotech, 450-33), 1X N2 (Gibco, 11520536) and 1X B27 (Gibco, 11500446).

*Generation of 2D organoids*

Prior to 2D culture, 3D organoids were pre-treated to promote proliferation and multipotency by culturing them in ENR-CV medium – ENR medium supplemented with 3 µM CHIR 99021 (Peprotech, 2520691) and 1 µM valproic acid (Sigma, P4543) for at least 3 days.
For 2D seeding, glass coverslips (13 mm diameter) were coated with extracellular matrix proteins. A drop of rat tail Collagen I (250 µg/mL Sigma; C3867) and Laminin 1 (100 µg/mL; Sigma, L2020) diluted in PBS was applied to each coverslip and incubated for 1h at 37°C. Excess matrix was removed, and coverslips was washed twice with PBS prior to use.
3D organoids were mechanically dissociated, typically using 4-5 wells per coverslip. Cells were centrifuged at 200 × g for 3 min at 4°C, and the pellet was resuspended in ENR-CV medium. A 50 µL drop of the cell suspension was added onto each coated coverslip. After 1h incubation at 37°C in a humidified incubator with 5% $CO_2$, 350 µL of ENR-CV medium supplemented with 10 µM ROCK inhibitor Y27632 (Santa Cruz) was added. Medium was changed daily. 2D organoids were maintained in ENR-CV + ROCKi for one day, then switched to differentiation medium containing 10 µM ROCKi. All experiments were performed 3-5 days after seeding.
To promote goblet cell differentiation, 3 µM DAPT (Tebu-bio, 28T6202) and IWP-2 (Tebu-bio, 28T2702) were added to fresh medium (*25*).
For blebbistatin treatment, 15 µM blebbistatin or vehicle (DMSO) was added to the 2D monolayers for 3h. Samples were then fixed and processed for analysis.

*Immunofluorescence on 2D organoids*

The organoid monolayers were fixed in 4% PFA for 10 min and washed 3 times with PBS. Samples were permeabilized with 0.1% Triton X-100 (Sigma) for 10 min, then washed 3 times with PBS. After 1h in the blocking solution (10% FBS/PBS), primary antibody was added (diluted in 10% FBS/PBS) and incubated O/N at 4°C. After three washes of 5 min each in PBS, secondary antibody was added with DAPI and with or without WGA and phalloidin, in 10% FBS/PBS and incubated 1h at RT. Samples were washed three times for 5min each and mounted on slides using mounting agent (Aqua-Poly/Mount, Polysciences). To differentiate goblet and Paneth cells, lysozyme staining was used and goblet cells were identified as lysozyme$^-$ and WGA$^+$ cells.

*Immunoblotting*

For each sample, small intestines were dissected and opened longitudinally. Tissue was incubated in solution containing 30 mM EDTA and 1 mM DTT in PBS for 20 min on ice, followed by 30 mM EDTA/PBS for 15 min at 37°C. The epithelium was detached from the mucosa by vigorous shaking and the epithelial fraction was collected by centrifugation and lysed in RIPA buffer. Protein concentration was quantified using DC protein assay (Bio-rad) and normalized, then 4X Laemmli buffer was added. 25 µg of total protein was used for electrophoresis on 4-20% sodium dodecyl sulfate (SDS) polyacrylamide gels under reducing conditions (Ready Gel, Bio-Rad), and transferred to a nitrocellulose membrane (Bio-Rad) following standard procedures. The membranes were blocked with blocking solution (Everyblot, Bio-Rad) for 30 min. The membranes were incubated with primary antibodies O/N at 4°C, followed by incubation with Horseradish peroxidase-conjugated secondary antibodies for 1 h at RT. The immunoreactive bands were visualized using chemiluminescence detection reagent (ECL Calirity, Bio-Rad) and ChemiDoc imaging system (Bio-Rad)

Confocal 3D imaging

To ensure consistency across samples, all image stacks were acquired at the mid-region of the villi. Imaging was performed using an inverted, laser-scanning confocal microscope LSM880 (Zeiss) equipped with 405 nm, Argon, 561 nm, and 633 nm lasers. A 63× oil immersion objective was used to acquire images at a resolution of 1024 × 1024 pixels. Z-stacks were collected with a step size of 0.3 µm.

*High resolution (Airyscan) 3D tissue imaging*

Images were acquired on laser-scanning confocal microscope LSM880 (Zeiss) equipped with an Airyscan module, using 63X oil objective, at a resolution of 1024x1024 pixels. Z-stacks were collected with a step size of 0.2 µm. Airyscan Z-stacks were processed in Zen software (Zeiss) using the Airyscan processing module.

*RIM*

The 3D images were acquired using an inverted microscope (TEi Nikon) equipped with a ×100 magnification, 1.49 NA objective (CFI SR APO 100XH ON 1.49 NIKON). A SCMOS camera (ORCA-Fusion, Hamamatsu) was used. A commercial acquisition software (INSCOPER SA) 3DRIM acquisition. A collimated beam with a diameter of 2.2 mm was generated using fast diode lasers (Oxxius) with wavelengths centered at 488 nm (LBX-488-200-CSB). The polarized beam was rotated at an angle of 5° before hitting an X10 beam expander (GBE04-A) and producing an 18 mm TEM00 beam through a spatial light modulator. As described previously (*51*), a fast spatial light phase binary modulator (QXGA fourth dimension) was conjugated to the image plane to generate 200 random illuminations through each plane. As described previously (*51*) and on GitHub (https://github.com/teamRIM/tutoRIM), 3D image reconstruction was then performed. We used a prefiltering Wiener parameter equal to 0.08, an inverting parameter equal to 0.08, and an equalization parameter equal to 0.1.

Image analysis

All images were processed for brightness and contrast adjustments and analyzed using Fiji software. Apical cell area was measured by segmentation of ZO-1 staining using the Tissue Analyzer plugin. Fracture area and junctional intensity were manually measured using the Fiji

Measurement Tool. Line profile intensity was also manually measured by tracing a line of consistent length and using the Plot Profile module. Average junctional E-cadherin signal intensity was plotted along the re-scaled junctional length in R. The cell shape index was calculated as (perimeter/$\sqrt{area}$) and measured manually based on the ZO-1 signal. Mucus volume measurement was performed using Imaris software, with 3D segmentation based on the WGA signal using the Cell function.

<u>Statistics</u>

In all figures, n represents the number of mice and/or independent organoid experiments. Statistical analyses were performed using GraphPad Prism 8.0.1. For each experiment, normality was assessed using the Shapiro-Wilk test. For normally distributed values, differences between two groups were analyzed using an unpaired t-test, while non-normally distributed values were analyzed using the Mann-Whitney test. For comparisons across three or more groups, multiple t-tests with FDR correction were used.

Data are presented as mean ± SD. Symbols used are: ns: $p > 0.05$; *: $p \leq 0.05$; **: $p \leq 0.01$; ***: $p \leq 0.001$; ****: $p \leq 0.0001$.

### Theory Note

In this Supplementary Theory Note, we provide details on the fracture mechanics of intestinal epithelia. In an intestinal epithelium, goblet cells swell and push the adjacent enterocytes. Such mechanical challenge strains the junctions between enterocytes that surround the goblet cell and cause small cracks. We first assume the crack formation can be completely determined by the mechanical interaction between the goblet cell and a single junction between neighboring enterocytes, and evaluate the detachment of cell-cell junction (Section 1). This model can well explain the fact that the swelling of goblet cells promotes crack formation, but fails to explain why a tissue with more compliant cells shows less cracks. Such inconsistency indicates that the crack formation might not solely be a result of localized mechanical interaction, but also be dependent on the mechanical property of the whole tissue. We then consider a multiscale tissue model that involves the swelling of goblet cells and formation of cracks at cell-cell junctions as a function of surrounding tissue rheology (Section 2).

#### *1. Crack mechanics of cell-cell interface*

A cell-cell interface mainly contains intercellular junctions, plasma membrane and cell cortices – thin layers of actomyosin meshwork beneath the plasma membrane. Cell–cell junctions are composed of transmembrane proteins that mediate adhesion between adjacent cell surfaces, which include both the plasma membrane and the underlying cortex. These surfaces form an elastic structure capable of withstanding mechanical forces from the environment (e.g., swollen goblet cells). As mucus granules accumulate, the internal pressure within goblet cells may increase and exceed that of neighboring enterocytes. This pressure imbalance generates a tensile force, $F$, along the goblet–enterocyte interface (see Fig. S3C for schematic). According to the Young–Laplace equation, this force $F$ is positively correlated with both the pressure difference and the size of the goblet cell.

Previous studies indicate that the tensile force $F$ could act as a "peeling force" and cause interface debonding (*22*, *52*, *53*). To check if goblet cells induce junctional cracks in a similar way, here we propose a mechanical model that considers the deformation and debonding of cell-cell interface. Given that most junctional fractures happen to tight junctions, which are located near the apical surface of the epithelium, it is reasonable to assume that the deformation and debonding of the cell-cell interface mainly happen in a two-dimensional plane (i.e., the tissue surface). At the tissue surface, the cell-cell interface can be depicted as a pair of cell boundaries that is initially linked with tight junctions and withstands peeling force $F$ at one end. Given the mechanical properties of relevant cellular components, the cell-cell interface can be modelled as two initially parallel elastic beams (i.e., the cell boundaries) with bridging forces from tight junctions (Fig. S3D). The cellular components aside from the cell interface (e.g., the apical or lateral surface) also hinder the formation and extension of junctional cracks, thus also contribute to the bridging forces in our model.

We introduce curvilinear coordinates $(s, \theta)$, where $s$ denotes the arclength along the deformed cell boundary (i.e., the elastic beam), and $\theta$ is the angle between the initial boundary profile (parallel to $x$-axis) and the tangent to its current profile (see Fig. S3D for schematic). The bending energy of the elastic beam is given as $\int_0^L \frac{\kappa}{2}\left(\frac{\mathrm{d}\theta}{\mathrm{d}s}\right)^2 \mathrm{d}s$, with $\kappa$ the bending stiffness of cell boundary and $L$

the total length (the cell boundary is considered to be inextensible). The binding potential per unit length $V$ is dependent on the deflection distance ($y$) vertical to the initial profile of cell boundary, that is $V = V(y)$, and the total binding potential should be $\int_0^L V(y)\,\mathrm{d}s$. Besides, geometric variables are mutually dependent and yield the relation $\frac{\mathrm{d}y}{\mathrm{d}s} = \sin\theta$, which works as a geometric constraint in the model. In the end, the total energy of one cell-cell interface can be written as

$$W = \int_0^L \frac{\kappa}{2}\left(\frac{\mathrm{d}\theta}{\mathrm{d}s}\right)^2 \mathrm{d}s + \int_0^L V(y)\,\mathrm{d}s + \int_0^L \gamma(s)\left(\frac{\mathrm{d}y}{\mathrm{d}s} - \sin\theta\right)\mathrm{d}s, \qquad (S1)$$

where $\gamma(s)$ serves as a Lagrange multiplier.

The local minimum of the total energy $W$ in Eq. (S1) corresponds to the equilibrium state of the cell-cell interface. The minimization of energy $W$ results in the following set of nonlinear equations that describe the current profile of cell-cell interface:

$$\begin{aligned} \kappa\frac{\mathrm{d}^2\theta}{\mathrm{d}s^2} + \gamma\cos\theta &= 0, \\ \frac{\mathrm{d}\gamma}{\mathrm{d}s} &= \frac{\partial V(y)}{\partial y}, \\ \frac{\mathrm{d}y}{\mathrm{d}s} &= \sin\theta. \end{aligned} \qquad (S2)$$

Specifically, the binding potential $V(y)$ has two sources: (i) the tight junction that tends to bind cell boundaries together, (ii) the cellular structures beneath the tissue surface that hinder the deflection of cell boundaries. The binding potential of the tight junction can be modeled as a van der Waals-like potential (Israelachvili, 1992): $V_{\mathrm{TJ}}(y) = w_{\mathrm{TJ}}\left[\left(\frac{d}{y}\right)^9 - \left(\frac{d}{y}\right)^3\right]$, with $w_{\mathrm{TJ}}$ a pre-factor that characterizes the adhesion strength, and $d$ an intrinsic interaction distance between the cell boundaries. Even without external mechanical perturbations (e.g., the swelling of goblet cells nearby), the pair of cell boundaries still has a small gap that separate two cells. The width of this gap (i.e., the homeostatic distance between cell boundaries) is $2d_0$ (see Fig. S3D for schematic, and we can easily obtain $d_0 = \sqrt[6]{3}d$ by considering the local minimum of $V_{\mathrm{TJ}}$). The binding potential that arises from external cell components is assumed to be $V_{\mathrm{ext}}(y) = \frac{w_{\mathrm{ext}}}{2}\left(\frac{y}{d_0} - 1\right)^2$, with $w_{\mathrm{ext}}$ a pre-factor that characterize the bridging strength. The external cell components would not affect the mechanical state of cell-cell interface in the homeostatic state of the tissue, but will impose a bridging force that gradually increase with the extension of junctional cracks. Overall, we have the binding potential $V(y) = V_{\mathrm{TJ}}(y) + V_{\mathrm{ext}}(y)$.

To solve the equation system in Eq. (S2), we still need to specify the conditions at two endpoints of the cell-cell interface (i.e., the elastic beams). One can notice the parameter $\gamma(s)$ acts as the shear force in the elastic beam. On the side away from the goblet cell (i.e., $s = 0$), there is neither bending moment (which is proportional to $\frac{d\theta}{ds}$) nor shear force ($\gamma$): $\frac{d\theta}{ds} = 0$ and $\gamma = 0$ at $s = 0$. At the side linked to the goblet cell (i.e., $s = L$), the goblet cell generates a peeling force $F$ vertical to the cell-cell interface. This external force acts as a shear force. Thus, we have $\frac{d\theta}{ds} = 0$ and $\gamma = F$ at $s = L$. To simplify the analysis, we non-dimensionalize the parameters. The length-related parameters are normalized by the homeostatic distance $d_0$, such that $\bar{y} = y/d_0$, $\bar{s} = s/d_0$, and $\bar{L} = L/d_0$. Forces are normalized by the adhesion strength $w_{\mathrm{TJ}}$, yielding $\bar{\gamma} = \gamma/w_{\mathrm{TJ}}$ and $\bar{F} = F/w_{\mathrm{TJ}}$. Under this formulation, the equilibrium configuration of the cell–cell interface depends only on two dimensionless parameters: (i) the effective bending rigidity $\bar{\kappa} = \kappa/(u_0^2 w_{\mathrm{TJ}})$, and (ii) the relative bridging strength $\bar{w} = w_{\mathrm{ext}}/w_{\mathrm{TJ}}$. Now Eq. (S2) becomes

$$\bar{\kappa}\frac{\mathrm{d}^2\theta}{\mathrm{d}\bar{s}^2} + \bar{\gamma}\cos\theta = 0,$$
$$\frac{\mathrm{d}\bar{\gamma}}{\mathrm{d}\bar{s}} = \sqrt{3}(\bar{y}^{-4} - \bar{y}^{-10}) + \bar{w}(\bar{y} - 1),$$
$$\frac{\mathrm{d}\bar{y}}{\mathrm{d}\bar{s}} = \sin\theta. \tag{S3}$$

To evaluate the extent of junctional cracking, we introduce the "crack opening displacement" (COD) as the deflection distance of the enterocyte boundary at its endpoint adjacent to the goblet cell, i.e., $y(s = L)$. In the initial regime, COD remains small and varies minimally with increasing peeling force $F$ (or the volume expansion of the goblet cell), indicating that the cell-cell junction still holds together. However, once the peeling force $F$ exceeds a threshold, COD rises sharply – or even discontinuously – indicating the onset and progression of a junctional crack, and the failure of the tight junction (Fig. S3E, F). These theoretical predictions are consistent with classical models (*23, 52–54*) and explain well the experimental observation that the swelling of goblet cells promotes junctional fractures (Fig. 2). Sensitivity analysis of key mechanical parameters further shows that junctional cracks can form more easily when the cell-cell interface is mechanically weakened (Fig. S3E, F). Specifically: (i) a decrease in the effective bending rigidity $\bar{\kappa}$ would lower the threshold for the crack nucleation, making the cell-cell interface more prone to breach (Fig. S3E); (ii) when external cell structures are impaired (characterized as smaller bridging strength $\bar{w}$), crack nucleation is only slightly affected, but subsequent crack propagation becomes easier (Fig. S3F). However, these predictions are inconsistent with the experimental results obtained upon depletion of myosin IIA (NMIIA): with fewer myosin molecules linking cortical actin filaments, the cell cortex and associated elastic structures become softer. In this scenario, junctional fractures are inhibited in experiments (Fig. 4E, F), rather than facilitated as predicted by the model (Fig. S3E, F).

In this model, the cell-cell interface is treated as an elastic structure. However, epithelial cells can actively remodel cell junctions and adjust their shapes and positions during homeostasis or in response to abnormal mechanical cues (*9*, *42*). To account for these, we next extend the model to include mechanical interaction at the tissue level, aiming to explore how the goblet cell-induced mechanical perturbation impact neighboring enterocytes and their junctions.

#### 2. *Crack mechanics of the whole tissue*

Epithelial cells interact mechanically through adhesive and elastic forces at the cell-cell interface (see also Section 1 for discussion). Such mechanical interactions between cells can be captured by multiscale tissue models such as vertex models. In classical vertex models (*39, 41*, *42*, *55–58*), each cell is described as a deformable polygon with multiple vertices, where each vertex represents a tricellular junction where cell boundaries meet, and on which the force balance is written. Although such models can well reproduce cell packing geometry (*42*), tissue rheology (*36, 41*) and related properties, they inherently assume that cell-cell interfaces remain straight and tightly connected – each cell interface being modeled as a straight line linking two vertices – and thus cannot capture junctional fractures.

Here, to investigate the formation mechanism of junctional cracks, we extend the classical vertex model by considering the cell-cell interface to be bendable and detachable (*23, 59*). To this end, we still model each cell-cell interface as two cell boundaries initially bonded together (as in Section 1), and discretize each cell boundary as a set of particles linked with elastic springs (Fig. S10A): the mechanical interactions among these particles endow the cell boundary with bending rigidity and extensibility, and the positions of these particles depict the profile of the cell boundary. Importantly, to mimic the adhesive effects of tight junctions, we endow the directly opposite particles (at the paired cell boundaries) with attractive forces to each other. In this way, this model can not only take the changes in cell shape and positions (as in classic vertex models) into account, but also allow for fractures of cell junctions. In the scenario without junctional fracture, this model gives similar predictions to the classic vertex models.

Analogous to the classical vertex models and the interface model proposed in Section 1, the mechanical energy of an enterocyte is mainly generated by changes in cell size (i.e., cell area in two-dimensional modeling) and elastic deformations of the cell surface/boundary (i.e., plasma membrane and cell cortex), the latter of which can be further divided into the boundary extension/contraction and the bending deformation. Thus, the cellular mechanical energy can be formulated as

$$U = \frac{K_A}{2}(A - A_0)^2 + \sum_{i=1}^{N} \frac{NK_L}{2}(L_i - L_0)^2 + \sum_{i=1}^{N} \frac{NK_B}{2}(\cos\theta_i - 1)^2 . \qquad (S4)$$

The first term of the energy $U$ comes from the change of the cell area, where $K_A$ is the area rigidity and $A$ (or $A_0$) is the current (or preferred) cell area. The second term is about the elastic extension/contraction of the cell boundary: let $K_L$ be the extension rigidity, and divide the whole boundary of one cell into $N$ pieces (with an equal length initially), whose current (or preferred) length is $L_i$ (or $L_0$). Correspondingly, the cell perimeter is $P = \sum_{i=1}^{N} L_i$ and the preferred perimeter is $P_0 = NL_0$. The last term represents the bending resistance of cell boundary, where $K_B$ is the bending rigidity and $\theta_i$ is the angle between neighboring pieces of the cell boundary. In numerical simulations, we set $N = 34$. Besides, as the single cells are initially in their round shapes (before being in contact with each other), we set a preferred cell radius $R = 20\mu\text{m}$ and thus calculate the preferred cell area as $A_0 = NR^2 \sin(2\pi/N)/2$.

The mechanical energy of a single cell can be non-dimensionalized as

$$u = \frac{U^{\mathrm{E}}}{K_A A_0^2} = \frac{1}{2}(a - 1)^2 + \sum_{i=1}^{N} \frac{k_l}{2}\left(l_i - \frac{p_0}{N}\right)^2 + \sum_{i=1}^{N} \frac{k_b}{2}(\cos\theta_i - 1)^2 , \qquad (S5)$$

with $a = A/A_0$ and $l_i = L_i/\sqrt{A_0}$ respectively the dimensionless area and length, and $k_l = NK_L/(K_A A_0)$ and $k_b = NK_B/(K_A A_0^2)$ respectively the dimensionless extensional rigidity and bending rigidity of the cell boundary. The key parameter in Eq. (S5) is the perimeter-to-area ratio $p_0 = P_0/\sqrt{A_0}$ (with $P_0 = NL_0$), which also referred to as "target shape index" in some references (*41*). This parameter determines the solid-fluid transition of epithelial tissues (*41*): the tissue is more of a solid when this shape index is below a threshold, and become fluid after the shape index exceeds the threshold. This is because cells with small perimeters are hard to change shape and position, whereas cells with large perimeters are floppy and tend to move under mechanical loading (*36*). Inspired by this, we also quantified the shape index of enterocytes to infer the mechanical properties of the tissue and individual cells. After depletion of myosin IIA (NMIIA), we found that the "shape index" increases (Fig 4J), indicating that some enterocytes become floppy and can more easily change their shape and position, while the tissue in general becomes more fluid.

In addition to the enterocytes discussed above, the intestinal epithelium also contains goblet cells, which are assumed to have the same mechanical/geometric parameters as enterocytes except for the "preferred cell area", denoted by $A_0^{\mathrm{G}}$. Following the same normalization procedure in Eq. (S5), one can obtain the dimensionless mechanical energy of a goblet cell as

$$u_{\mathrm{G}} = \frac{1}{2}\left(a - a_0^{\mathrm{G}}\right)^2 + \sum_{i=1}^{N} \frac{k_l}{2}\left(l_i - \frac{p_0}{N}\right)^2 + \sum_{i=1}^{N} \frac{k_b}{2}(\cos\theta_i - 1)^2,$$

which includes an additional geometric parameter: the normalized preferred area of goblet cells $a_0^{\mathrm{G}} = A_0^{\mathrm{G}}/a_0$. Thus, the total energy of the intestinal tissue can be written as

$$u_{\mathrm{t}} = \frac{1}{2}\sum_{m=1}^{N_{\mathrm{c}}}(a_m - a_0)^2 + \frac{k_l}{2}\sum_{m=1}^{N_{\mathrm{c}}}\sum_{i=1}^{N}\left(l_{m,i} - \frac{p_0}{N}\right)^2 + \frac{k_b}{2}\sum_{m=1}^{N_{\mathrm{c}}}\sum_{i=1}^{N}\left(\cos\theta_{m,i} - 1\right)^2, \qquad (S6)$$

where the value of $a_0$ can be set as 1 (for enterocytes) or $a_0^{\mathrm{G}}$ (for goblet cells). In our simulations, we first set $a_{\mathrm{G}0} = 0.2$ in the initial step, then increase its value to mimic the swelling process of goblet cells.

The movement of discrete particles at the cell boundaries is driven by the changes in the cell energy and cell-cell interactions. In our simulations, the spatio-temporal evolution of particle $\boldsymbol{r}_i$ of cell $m$ yields

$$\gamma \frac{\mathrm{d}\boldsymbol{r}_i}{\mathrm{d}t} = -\frac{\partial U_m}{\partial \boldsymbol{r}_i} + F_{m,i}^{\mathrm{int}}, \qquad (S7)$$

where $\gamma$ is the damping coefficient, $U_m$ is the mechanical energy of cell $m$ (see Eq. (S4) for definition) and $F_{m,i}^{\mathrm{int}}$ represents the force arising from the cell–cell interactions. To avoid overlapping between cells, we also introduce repulsive forces between discrete particles that belonging to different cells: we set the repulsive force as $f_0\left(-\frac{1}{\left|\boldsymbol{r}_{m,i}-\boldsymbol{r}_{n,j}\right|^2} + \frac{1}{{r_{\mathrm{e}}}^2}\right)\frac{\boldsymbol{r}_{m,i}-\boldsymbol{r}_{n,j}}{\left|\boldsymbol{r}_{m,i}-\boldsymbol{r}_{n,j}\right|}$, where $\boldsymbol{r}_{m,i}$ denotes the vertex $i$ of cell $m$, $f_0$ represents the intensity of repulsion, and $r_{\mathrm{e}}$ is an effective distance.

Supplementary Table 1. Key parameters used in the model

| Physical meaning | Symbol | Value | Reference |
|---|---|---|---|
| area rigidity | $K_A$ | $1 \times 10^7$ N/m$^3$ | (*56*) |
| extension rigidity | $K_L$ | $1 \times 10^{-4}$ N/m | (*56*) |
| bending rigidity | $K_B$ | $6.25 \times 10^{-125}$ J | (*59*) |
| damping coefficient | $\gamma$ | 0.1 Ns/m | (*56*) |
| repulsive strength | $f_0$ | $5\times 10^{-4}$ N/m | (*60*) |
| effective distance | $r_{\mathrm{e}}$ | 1.25 μm | (*61*) |

This model predicts that the nucleation and extension of junctional cracks depend not only on the size of goblet cells (characterized as $a_0^{\mathrm{G}}$), whose propagation would push the adjacent enterocytes and peel apart their cell junctions (in agreement with the experimental observation in Fig. 2 and the theoretical prediction in Section 1), but also depends on the general rheological property of the epithelia (implied by the cell shape index $p_0$): solid-like tissues are hard to deform under mechanical perturbations, and thus more likely to accumulate local stresses that can induce crack nucleation; in contrast, in fluid-like tissues, their enterocytes are floppy and can easily rearrange their positions and adjust their shape/size to adapt to abnormal mechanical forces. Such mechanical self-adjustment prevents stress accumulation and thus effectively protects the tissue from physical damage even if individual cells are mechanically vulnerable. These predictions can well explain the perturbation experiments on cell mechanics (Fig. 4): after depletion of myosin IIA, the epithelium becomes more fluidic (as the cellular shape index increases, see quantification in Fig. 4 J) and the cracks become fewer and smaller (Fig. 4F, L).

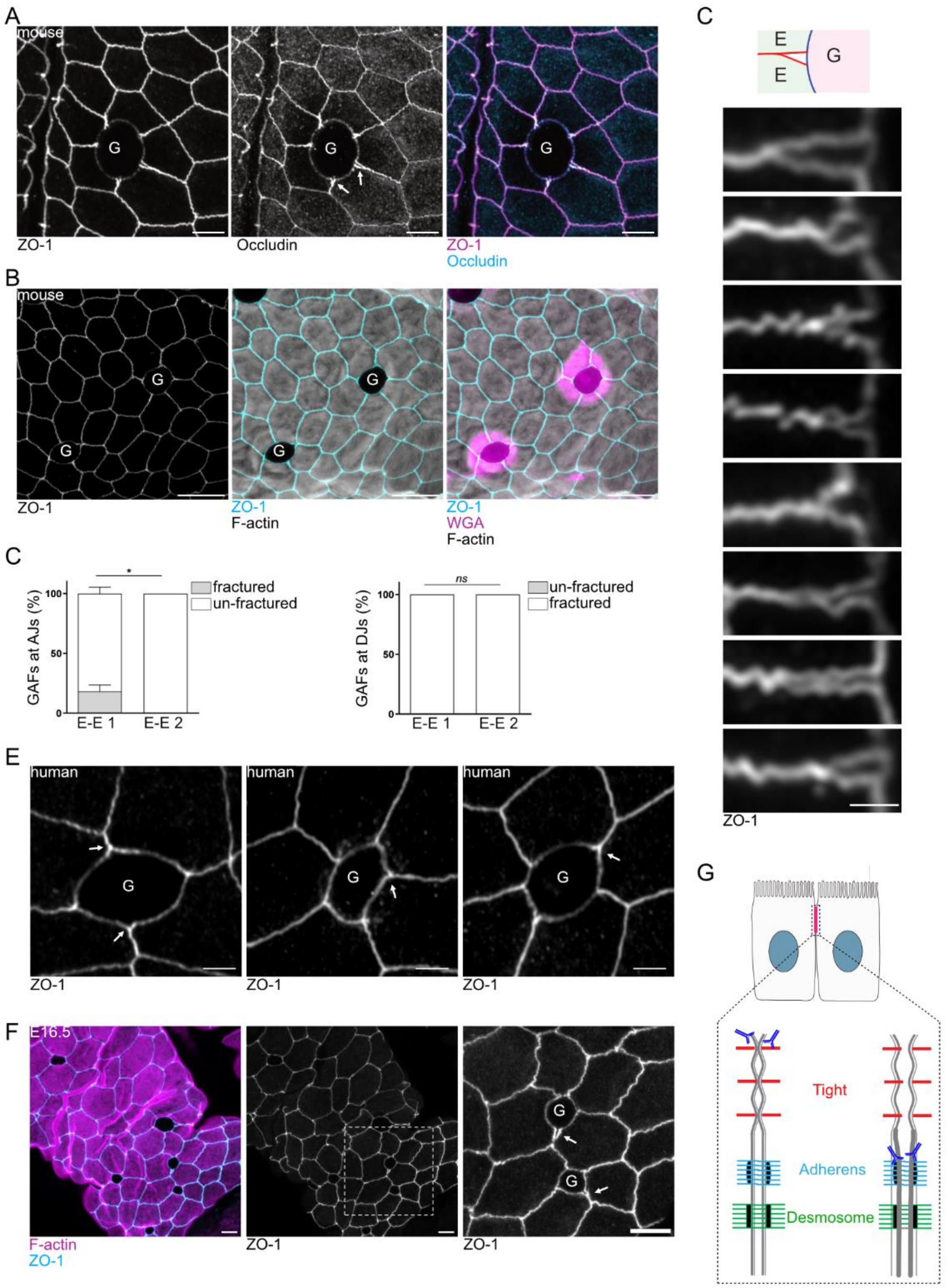


**Fig. S1.** (**A**) Representative *en face* images of murine tissue; maximum intensity projection (Z range, 3.5 µm). White arrows are showing fractures in 1st neighbor E-E junctions, Scale bars, 5 µm. (**B**) Representative *en face* images of murine intestinal tissue; maximum intensity projection (Z range, 3.2 µm). Scale bars, 10 µm. (**C**) Montage of high-resolution images showing fractures at the goblet-enterocyte-enterocyte tricellular junction. Scale bar, 2 µm. (**D**) Stacked bar graph showing average percentage of enterocytes associated with fractures at AJs (left) and desmosome level (right). Multiple t-test, *: $p < 0.05$, ns: non-significant; n=3 experiments. (**E**) Representative *en face* images of human ileum; maximum intensity projection (Z range, 1.4 – 1.8 µm). White arrows are showing fractures in 1st neighbor E-E junctions. Scale bars, 2 µm. (**F**) Representative images of E16.5 murine intestine. White arrows are showing GAFs in embryonic intestine; n=4 embryos. Scale bars, 5 µm. (**G**) Schematic representation of accessible E-cadherin staining.

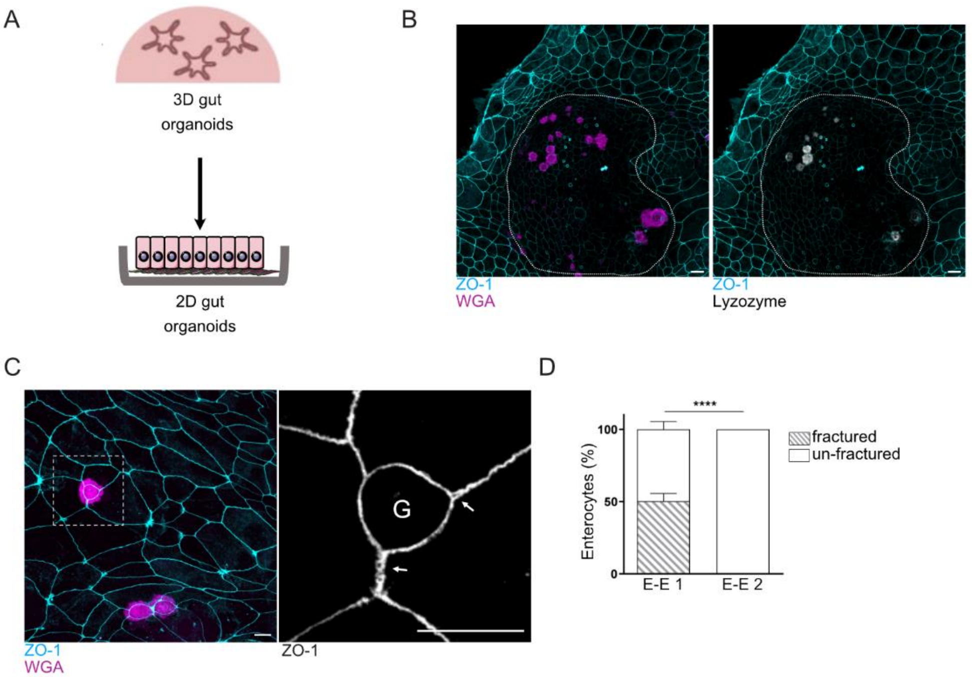


**Fig. S2.** (**A**) Schematic representation displaying the generation of 2D organoids from 3D organoids. (**B**) Representative images of 2D organoids stained for mucus with WGA (magenta), ZO-1 (cyan) and lysozyme (grey) showing the crypt-like (encircled area) and villus-like organization of 2D organoids; maximum intensity projection (Z range, 9.52 µm). Scale bars, 10 µm. GCs are identified as WGA+ Lyz- cells. (**C**) Representative images of 2D organoids stained for WGA (magenta), ZO-1 (cyan). White arrows are showing GC-associated fractures *in vitro*; maximum intensity projection (Z range, 3.8 µm). Scale bar, 10 µm. Boxed region is shown in higher magnification. Scale bar, 5 µm. (**D**) Stacked bar graph displaying average percentage of fractured enterocytes (GAFs). Multiple t-test, ****: $p < 0.0001$; n=4 experiments.

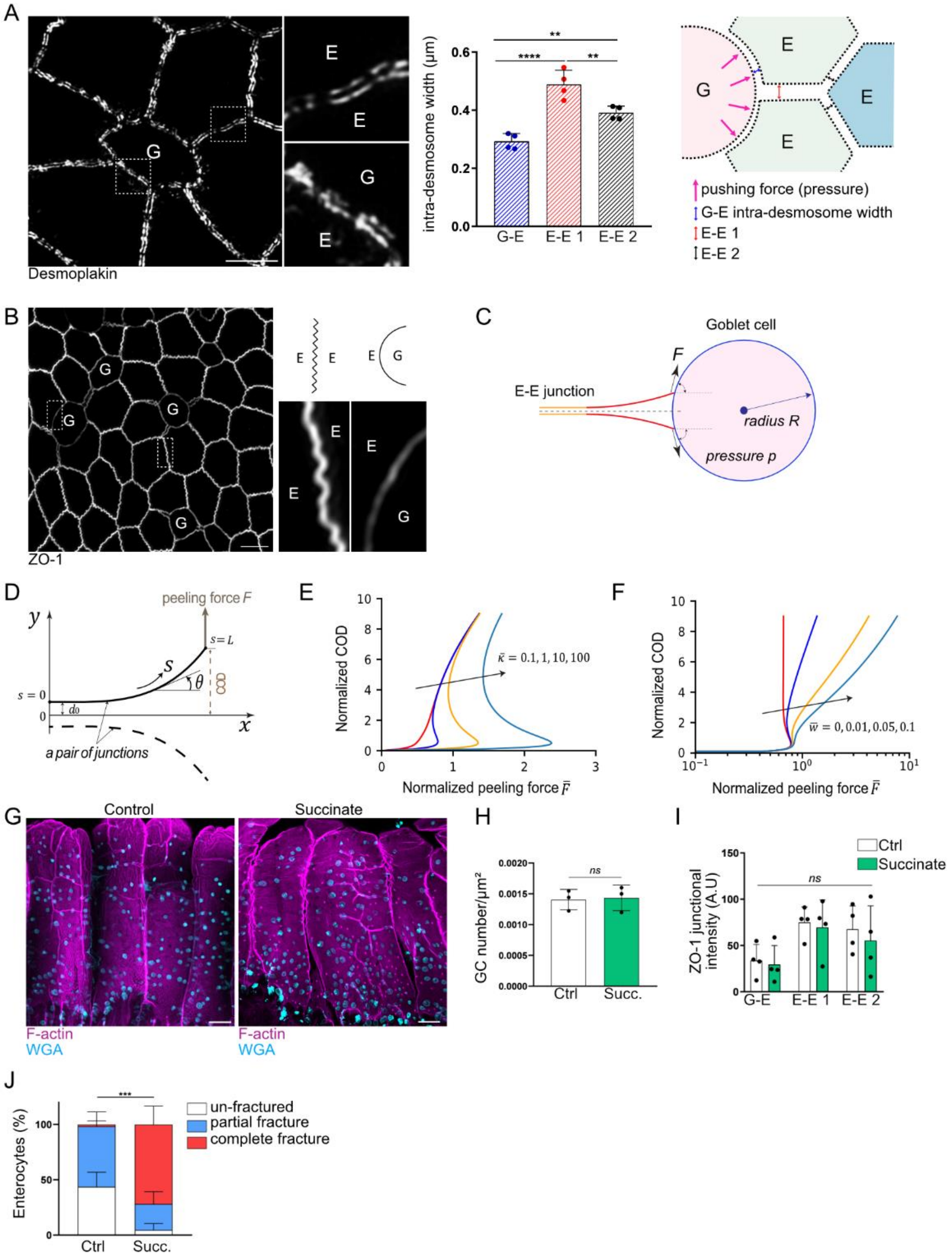


**Fig. S3. (A)** Left: super-resolution (RIM) *en face* images of intestinal tissue; boxed regions are shown in higher magnification; maximum intensity projection (Z range, 1.2 µm). Middle: Bar chart displaying average intra-desmosomal width. Right: schematic representation displaying intra-desmosomal width analysis. Multiple t-test, **: p < 0.01, ****: p < 0.0001; n=4 independent experiment. Scale bar, 2 µm. **(B)** Representative images showing E-E and G-E TJs; boxed regions are shown in higher magnification; maximum intensity projection (Z range, 3.1 µm). Scale bar, 5 µm. **(C)** Schematic to show the fracture mechanics of an E-E junction driven by forces exerted by the GC. **(D)** Schematic and parameter definitions of the crack mechanics model of cell-cell interface. **(E-F)** Sensitivity analysis of mechanical parameters to understand how junctional cracks develop due to the peeling force arising from the swelling of goblet cells (see also panel C). The crack growth is characterized as the normalized "crack opening displacement" (COD). Specifically, panels E and F analyze the influence of **(E)** the effective bending rigidity of the cell boundary ($\bar{\kappa}$, with $\bar{w} = 0.01$) and **(F)** the bridging strength of external components ($\bar{w}$, with $\bar{\kappa} = 1$), respectively. **(G)** Representative low magnification images showing whole villi of control and succinate treated tissue stained; maximum intensity projection (Z range, 13.95 µm and 11.2 µm). Scale bars, 50 µm. **(H)** Bar chart displaying average GC number per $\mu m^2$. Mann-Whitney test, ns: non-significant; n=3 experiments. **(I)** Bar charts showing average ZO-1 junctional intensity. Multiple t-test, ns: non-significant; n=4 experiments. **(J)** Stacked bar graph showing average percentage of un-fractured, partially or totally fractured GC-1st neighbor E-E junctions TJs. Multiple t-test, ***: p<0.001; n=4 experiments. Ctrl, control; Succ., succinate.

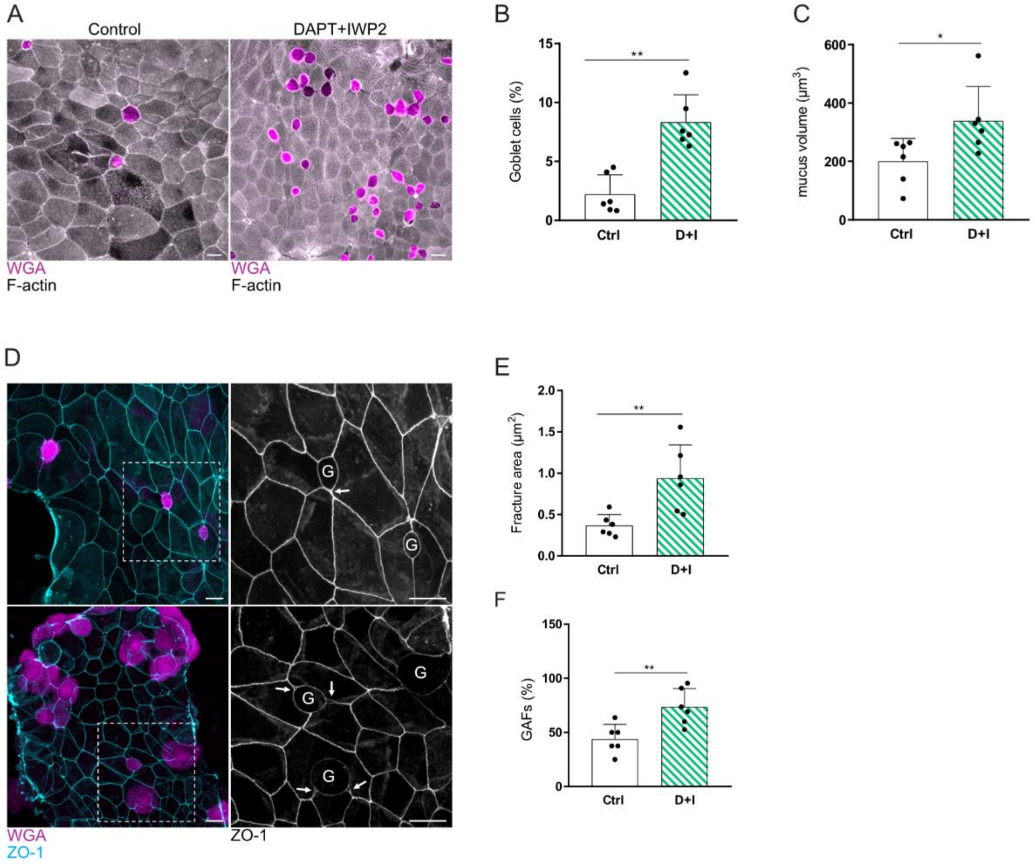


**Fig. S4.** (**A**) Representative images of 2D organoids treated or not with DAPT+IWP2, showing GC hyperplasia; maximum intensity projection (Z range, 3.3 µm and 8.4 µm). Scale bars, 10 µm. (**B**) Bar chart showing average percentage of goblet cell in 2D organoids according the conditions. Mann-Whitney test, **: $p < 0.01$; n=6 experiments. (**C**) Bar chart displaying mucus volume. Mann-Whitney test, *: $p < 0.05$; n=6 experiments. (**D**) Representative images of control and DAPT+IWP2-treated 2D organoids. Boxed regions are shown in higher magnification. White arrows are showing GAFs. Maximum intensity projection (Z range, 3.3 µm and 8.5 µm). Scale bars, 10 µm. (**E-F**) Bar charts showing average fracture area (**E**) and average percentage of GAFs at TJ level (**F**). Mann-Whitney test, **: $p < 0.01$; n=6 experiments. Ctrl, Control; D, DAPT; I, IWP2

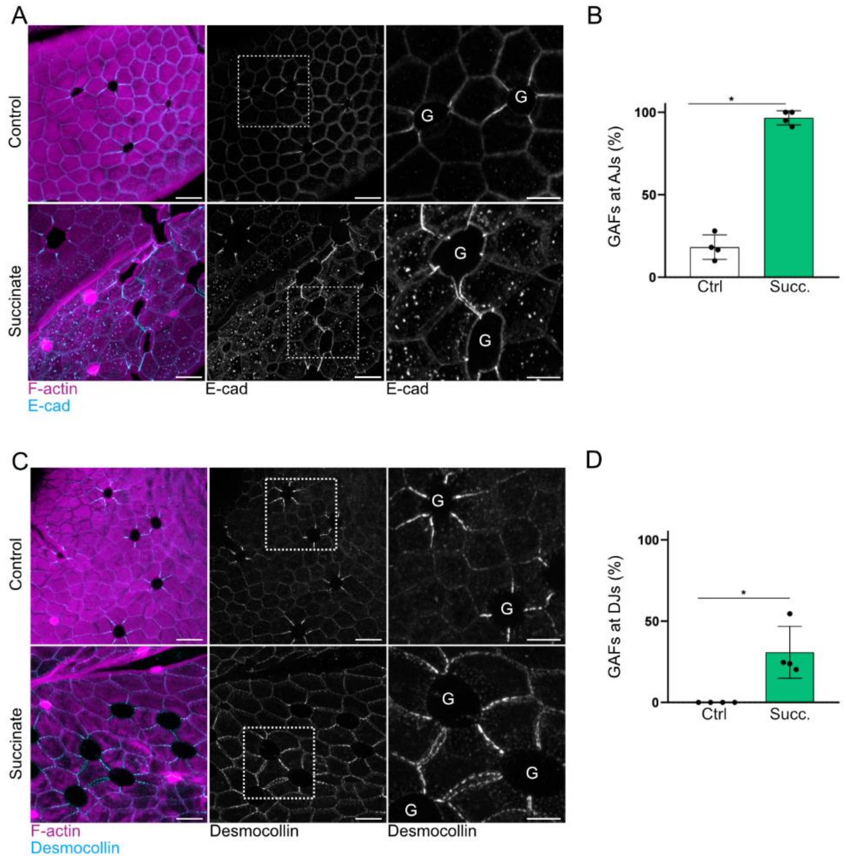


**Fig. S5.** (**A**) Representative images from control and succinate-treated mice; maximum intensity projection (Z range, 1.8 µm, 2.1 µm); scale bars, 10 µm. Boxed regions are shown in higher magnification. Scale bars, 5µm. (**B**) Bar chart displaying average percentage of GAFs at AJs level. Mann-Whitney test *: $p < 0.05$; n=4 experiments. (**C**) Representative images from control and succinate-treated mice.; maximum intensity projection (Z range, 4.2 µm and 3.3 µm); scale bars, 10 µm. Boxed regions are shown in higher magnification; scale bars, 5 µm. (**D**) Bar chart displaying average percentage of GAFs at desmosome level. Mann-Whitney test, *: $p < 0.05$; n=4 experiments. Ctrl, control; Succ., succinate.

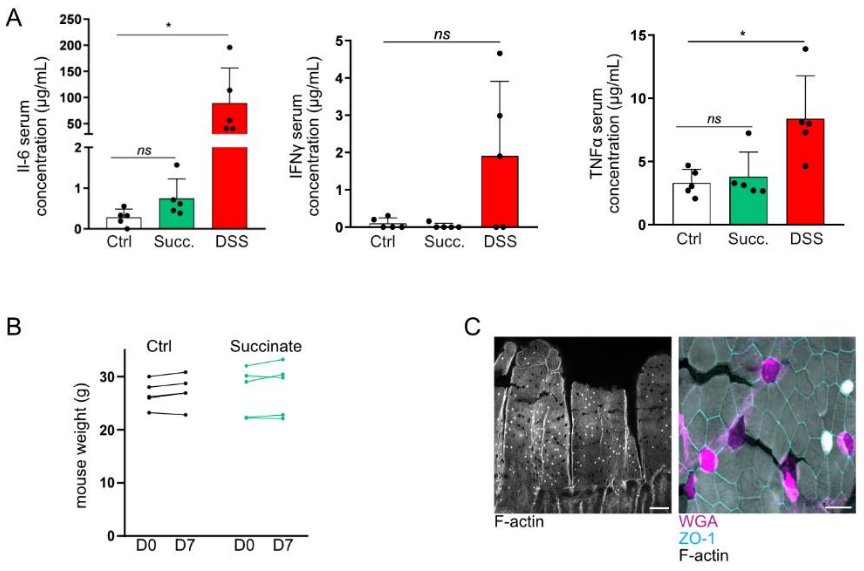


**Fig. S6.** (**A**) Bar charts showing IL-6, IFNγ and TNF-α average serum concentration. Multiple t-test, *: ns: non-significant, $p<0.05$; n=5 experiments. (**B**) Control and succinate mice weight plot. Multiple t-test; n=5 experiments. (**C**) Representative images of tissue from succinate-treated mice, treated with luminal shear stress *ex vivo.* Left: low magnification image, showing damage at the tip of the villi. Scale bar, 50 µm. Right: High-resolution image showing long-range fracturing emanating from GCs. Maximum intensity projection (Z range, 6.6 µm). Scale bar, 10 µm. Ctrl, Control; Succ., succinate.

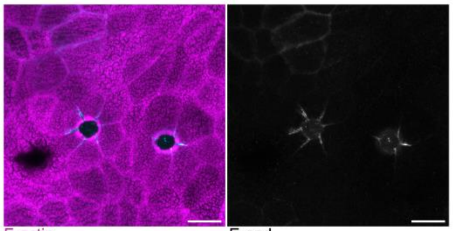


**Fig. S7**. Representative images of 2D organoid monolayers; maximum intensity projection (Z range, 2.7 µm). Scale bars, 10 µm.

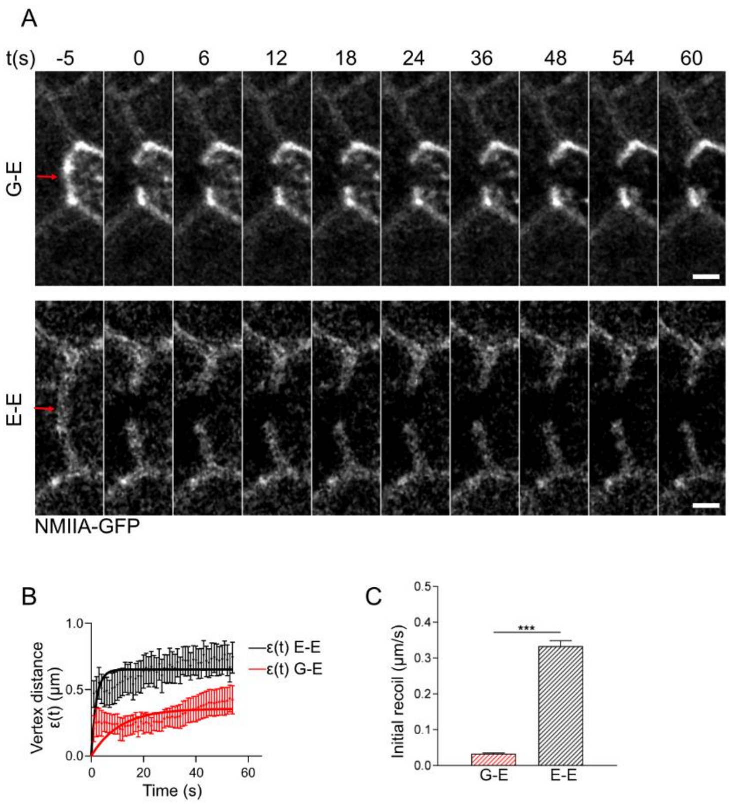


**Fig. S8.** (**A**) Representative montages showing junctional recoil of G-E (top panel) and E-E (bottom panel) AJs after laser cuts; NMIIA-GFP tissue. Scale bars, 2µm. (**B**) Vertex distance ε(t) plot in function of time for G-E (red) and E-E (black) AJs. (**C**) Bar chart showing average initial recoil speed for G-E and E-E AJs. Mann-Whitney test, ***: p<0.001; n=3 experiments.

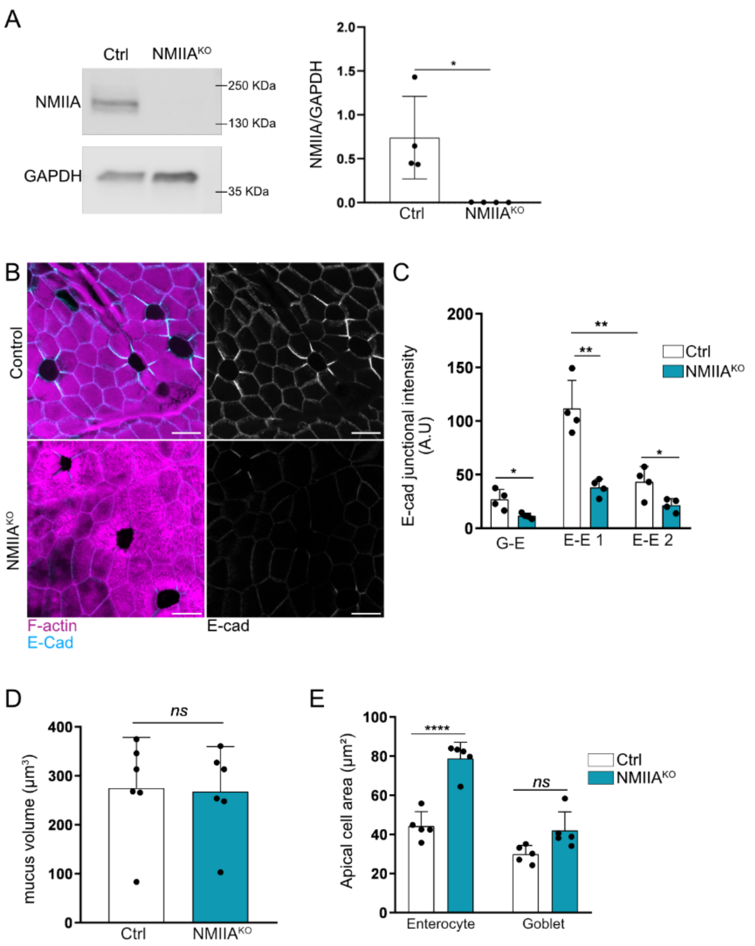


**Fig. S9**. (**A**) Right: Representative western blot for NMIIA from control and NMIIA$^{KO}$ small intestinal epithelium; GAPDH was used as loading control. Left: Bar chart showing NMIIA signal intensity normalized to GAPDH. Mann-Whitney test, *: p<0.05; n= 4 experiments. (**B**) Representative *en face* images of control and NMIIA$^{KO}$ tissue; maximum intensity projection (Z range, 2.4 µm and 1.8 µm). Scale bars, 10 µm. (**C**) Bar chart showing average E-cadherin junctional intensity in control or NMIIA$^{KO}$ tissue. Multiple t-test, *: p < 0.05, **: p<0.01; n=4 independent experiments. (**D**) Bar graph displaying average mucus volume. Mann-Whitney test, ns=non-significant, p>0.05; n=5 independent experiments. (**E**) Bar chart displaying average apical cell area. Multiple t-test, ns: non-significant; ****: p < 0.0001; n=5 experiments.

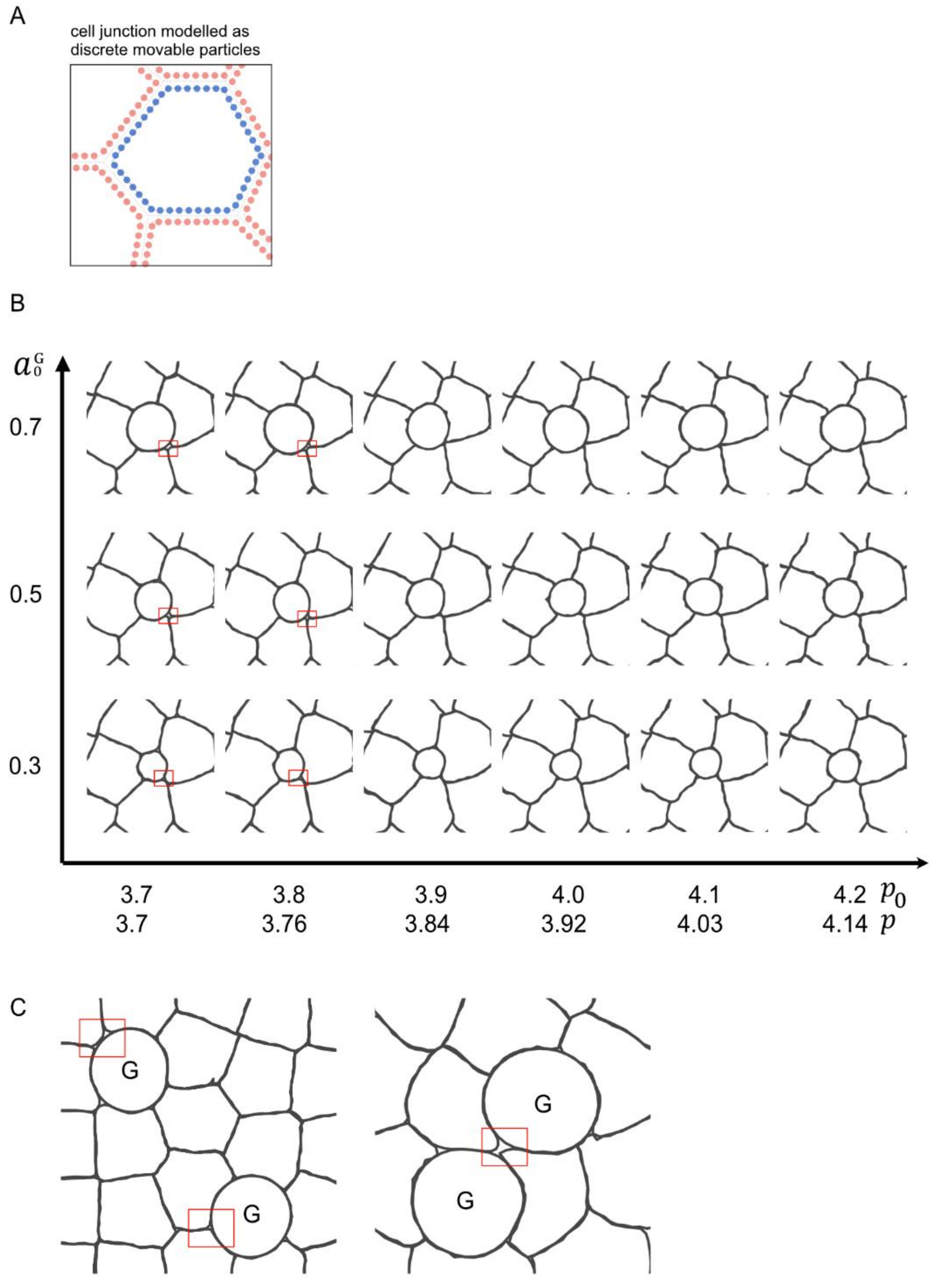


**Fig. S10.** (**A**) Cartoon representation of the tissue mechanics model, which treats the cell boundaries as discrete movable particles to track the cellular shape change and crack formation at cell-cell junctions. (**B**) Simulations of the intestinal epithelial junctions based on the tissue mechanics model (see Theory Note for details). The two key parameters are the preferred area ($a_0^G$, see $y$-axis) of the goblet cell (which has otherwise identical properties compared to the other surrounding cells), and the preferred shape index of all cells in the tissue ($p_0$, see $x$-axis). (**C**) Model simulation showing cumulative positional effect of GCs on fracture area. Goblet cells shown are either separated by 2 rows (left) of 1 row of enterocytes (right); $a_0^G$ = 0.7, $p_0$= 3.8.

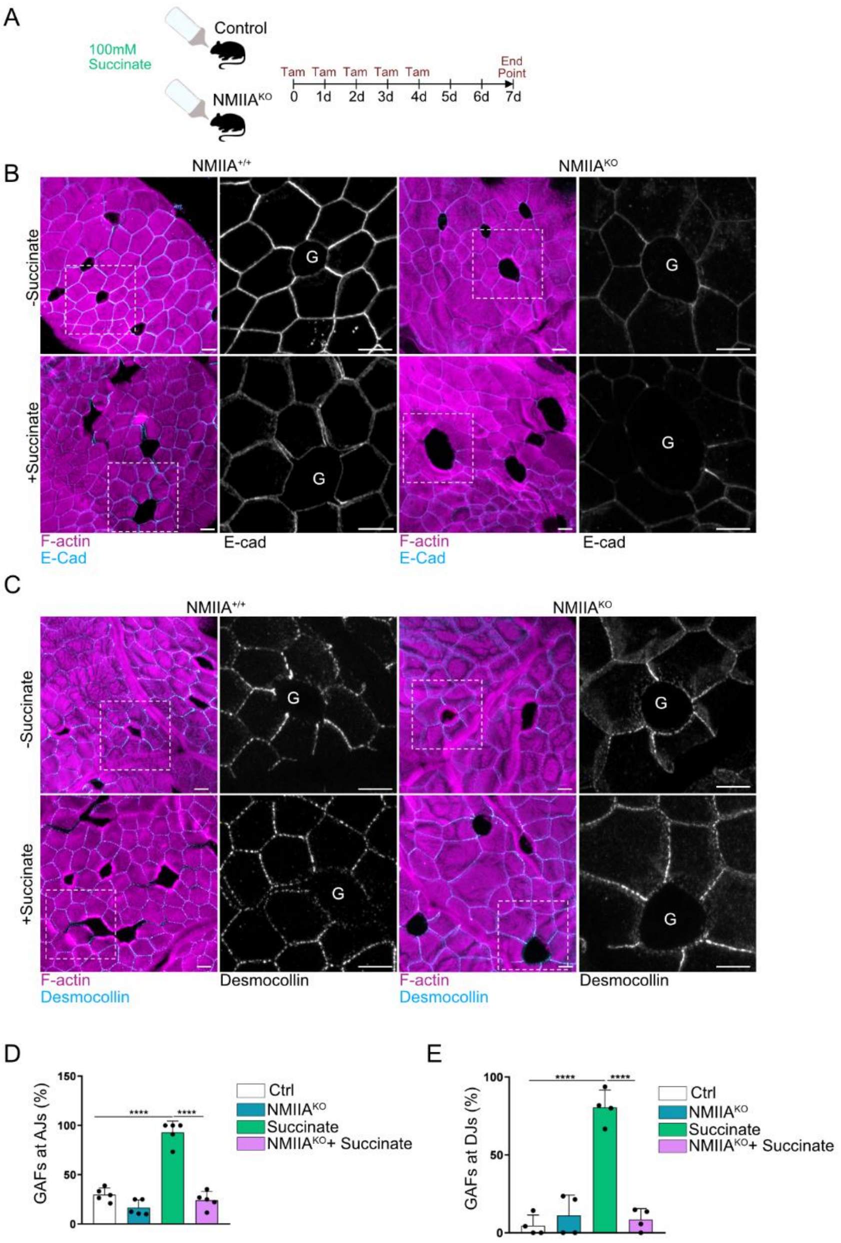


**Fig. S11**. (**A**) Schematic representation of succinate treatments in control (NMIIA$^{+/+}$) or NMIIA$^{KO}$ mice. (**B**) Representative images of tissues from NMIIA$^{+/+}$ and NMIIA$^{KO}$ mice with and without succinate treatment; maximum intensity projection (Z range, 2.4 – 4.2 µm). Boxed regions are shown in higher magnification. Scale bars, 10 µm. (**C**) Representative images from NMIIA$^{+/+}$ and NMIIA$^{KO}$ mice with and without succinate treatment; maximum intensity projection (Z range, 1.4 – 2.8 µm). Boxed regions are shown in higher magnification. Scale bars, 10 µm. (**D**) Bar chart displaying average percentage of GAFs at AJ level. Multiple t-test, ****: $p < 0.0001$; n=5 experiments. (**E**) Bar chart showing average percentage of GAFs at the desmosome level. Multiple t-test, ****: $p < 0.0001$; n=4 experiments.

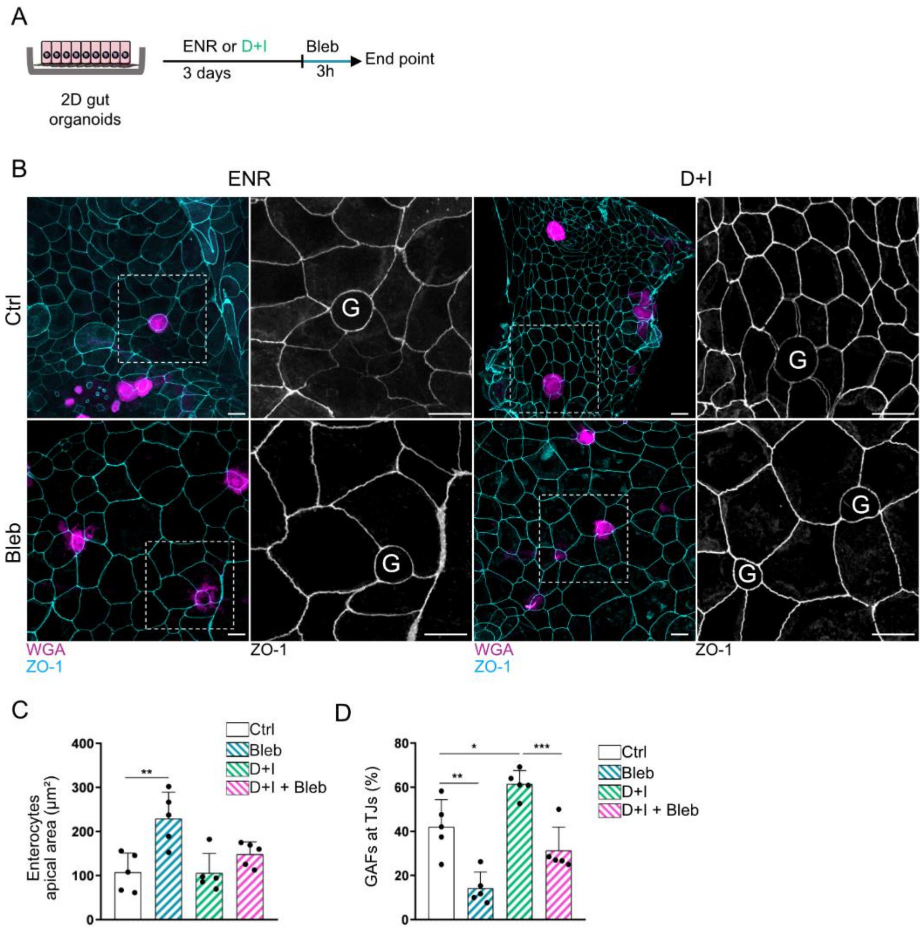


**Fig. S12**. (**A**) Schematic representation of blebbistatin treatment in 2D organoids pre-treated with DAPT (D) and IWP2 (I). (**B**) Representative images of 2D organoids treated with blebbistatin (Bleb) or vehicle control (DMSO). Scale bars, 10 µm. (**C**) Bar chart displaying enterocytes' average apical cell area according to the culture condition. Multiple t-test, **: p<0.01. (**D**) Bar graph showing the percentage of GAFs at the TJ level. Ctrl, vehicle control (DMSO); Multiple t-test, *: p<0.05, **: p<0.01; ***: p<0.001. D, DAPT; I, IWP2.

**Supplementary Materials – References**
